# Codependent excitatory and inhibitory plasticity accounts for quick, stable and long-lasting memories in biological networks

**DOI:** 10.1101/2021.04.01.437962

**Authors:** Everton J. Agnes, Tim P. Vogels

## Abstract

The brain’s functionality is developed and maintained through synaptic plasticity. As synapses undergo plasticity they also affect each other. The nature of such “codependency” is difficult to disentangle experimentally, because multiple synapses must be monitored simultaneously. To help understand the experimentally observed phenomena, we introduce a framework that formalises synaptic codependency between different connection types. The resulting model explains how inhibition can gate excitatory plasticity, while neighbouring excitatory-excitatory interactions determine the strength of long-term potentiation. Furthermore, we show how the interplay between excitatory and inhibitory synapses can account for the quick rise and long-term stability of a variety of synaptic weight profiles, such as orientation tuning and dendritic clustering of co-active synapses. In recurrent neuronal networks, codependent plasticity produces rich and stable motor cortex-like dynamics with high input sensitivity. Our results suggest an essential role for the neighbourly synaptic interaction during learning, connecting micro-level physiology with network-wide phenomena.

---

The remarkable property of synapses to change – synaptic plasticity – is thought to be the brain’s fundamental mechanism for learning, and it has generated a large body of theoretical and experimental work^1–4^. Based on Hebb’s postulate and early experimental data, theories have focused on the idea that synapses change solely based on the activity of their pre- and postsynaptic counterparts^5–12^, defining synaptic plasticity as predominantly a local process that is controlled by global modulatory signals. However, experimental evidence^13–22^ has pointed towards learning mechanisms that act non-locally at the mesoscale, taking into account the activity of multiple synapses and synapse types nearby. For example, excitatory synaptic plasticity (ESP) has long been known to rely on intersynaptic cooperativity by way of elevated calcium concentrations from the activation of multiple presynaptic excitatory synapses^17–20^. Interestingly, GABAergic, inhibitory synaptic plasticity (ISP) has also been shown to depend on the activation of neighbouring excitatory synapses: ISP is blocked when excitatory synapses are deactivated^13,14^, and the magnitude of the changes depends on the ratio of local excitatory and inhibitory currents (EI balance)^13^. Finally, inhibitory currents can affect ESP, either flipping the direction of efficacy changes according to the absence or presence of inhibitory currents in the vicinity of the synapse^15,16^, or maximising ESP during local disinhibition^23^ caused by modulatory inputs^24^. Longterm potentiation (LTP) at excitatory synapses has also been shown to depend on the distance and timing between successive LTP inductions of neighbouring excitatory synapses^25^. Moreover, Hebbian LTP has been shown to trigger long-term depression (LTD) at neighbouring synapses^21^ through a heterosynaptic plasticity mechanism. There is currently no unifying framework to incorporate these experimentally observed inter-dependencies at the mesoscopic level of synaptic plasticity. Existing models typically aim to explain, e.g., how cell assemblies are formed and maintained^26,27^. In these studies, local plasticity rules are typically complemented with non-local processes such as normalisation of excitatory synapses^27^, or modulation of inhibitory synaptic plasticity by the average network activity^10^, for stability. Moreover, intricate spatiotemporal dynamics, such as the activity patterns observed in motor cortex during reaching movements^28^, can only be reproduced when inhibitory connections are optimised (i.e., hand tuned) by iteratively changing the eigenvalues of the connectivity matrix towards stable values^29^, or learned by non-local supervised algorithms such as FORCE^30,31^. However, models that rely on connectivity changes triggered by non-local quantities are usually based on the optimisation of network dynamics^29–31^, and often don’t reflect biologically relevant mechanisms (but see Kirchner and Gjorgjieva ^32^).

In order to fill the theoretical gap in non-local, mesoscopic synaptic plasticity rules, we introduce a new model of *codependent* synaptic plasticity that takes into account the direct interaction between different neighbouring synapses. Our model can account for a wide range of experimental data on excitatory plasticity and receptive field plasticity of excitatory and inhibitory synapses, and makes predictions for future experiments involving multiple synaptic stimulation. Furthermore, it provides a mechanistic explanation for experimentally observed synaptic clustering and for how dendritic morphology plays a role to facilitate the emergence of single (clustered) or mixed (scattered) feature selectivity. Finally, we show how naïve recurrent networks can grow into strongly connected, stable and input sensitive circuits showing amplifying dynamics.

## Results

We developed a general theoretical framework for synaptic plasticity rules which accounts for the interplay between different synapse types during learning. In our framework, excitatory and inhibitory synapses change according to the functions *ϕ*_E_(*E, I*; *PRE, POST*) and *ϕ*_I_(*E, I*; *PRE, POST*), respectively (Fig. 1A). The signature of the codependency between neighbouring synapses is given by *E* and *I*, which describe the recent postsynaptic activation of nearby excitatory and inhibitory synapses, respectively. The activity of the synapse’s own pre- and postsynaptic neurons, i.e., the synapse’s local activity, is described by the variables *PRE* and *POST*, respectively.

**FIG. 1.**
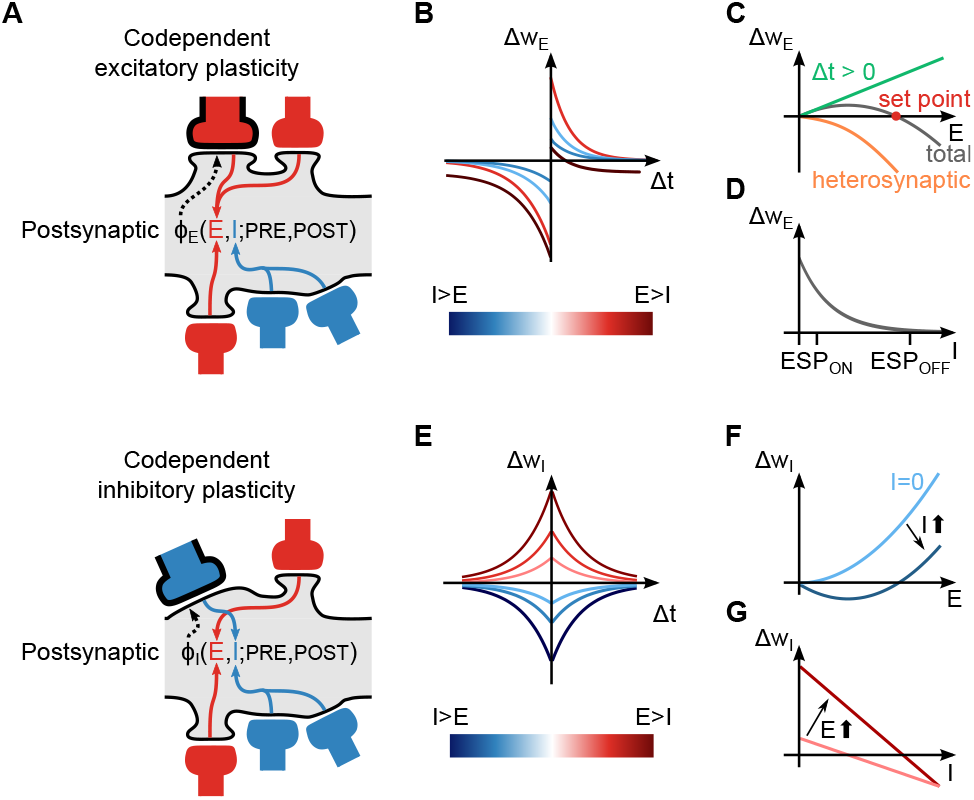
Codependent synaptic plasticity model. **A**, Codependent excitatory (top) and inhibitory (bottom) plasticity. Plasticity of a synapse (highlighted with black contour) depends on the activation of neighbouring excitatory (red) and inhibitory (blue) synapses, together with local pre- and postsynaptic activity. **B**, Excitatory weight change, Δ*w*_E_, is a function of the time interval between post and presynaptic spikes, Δ*t*, and neighbouring synaptic inputs, *E* and *I*. **C**, Excitatory inputs, *E*, control Hebbian-LTP (green line; Δ*t >* 0) and heterosynaptic plasticity (orange line), which combined (grey line) create a common set point for the total excitatory input (red dot). **D**, Inhibitory inputs, *I*, gate excitatory plasticity (‘ESP_ON_’ *vs* ‘ESP_OFF_’). **E**, Inhibitory weight change, Δ*w*_I_, is a function of Δ*t* and neighbouring synaptic inputs (as in panel B). **F** and **G**, Synaptic changes in inhibitory synapses as a function of excitatory (panel F) and inhibitory (panel G) inputs.

We modelled *E* and *I* as variables that depend on neighbouring synaptic currents: calcium influx through N-methyl-D-aspartate (NMDA) channels for *E*, and chloride influx through *γ*-Aminobutyric acid type A (GABA_A_) channels for *I* (see Methods). The impementation of excitatory and inhibitory plasticity rules vary slightly, as follows below.

### Codependent excitatory plasticity model

The rule *ϕ*_E_(*E, I*; *PRE, POST*) by which excitatory synaptic efficacies change is constructed similarly to classic spike-timing dependent plasticity (STDP) models^17,33^: pre-before-post spike patterns elicit potentiation while post-before-pre elicits depression (Fig. 1B). In addition, synaptic changes are modulated by neighbouring excitatory and inhibitory activity such that the learning rate for potentiation increases linearly with the magnitude of neighbouring excitatory inputs^17,18,20^ (Fig. 1C, green line). This potentially de-stabilising positive feedback, in which potentiation leads to bigger excitatory currents which in turn leads to more potentiation, is counterbalanced by including an experimentally observed heterosynaptic term^10^ that weakens synapses via a quadratic dependency on the neighbouring excitatory currents^21,34^ (Fig. 1C, orange line). Together, pre-before-post potentiation and and heterosynaptic weakening form a fixed-point in the dynamics of synaptic weights. Weak to intermediate excitatory currents elicit synaptic strengthening. Strong currents induce synaptic weakening (Fig. 1C, grey line). In addition to neighbouring excitatory-excitatory effects, we constructed the model such that elevated inhibition blocks excitatory plasticity^35^: Only when synapses are disinhibited can excitatory plasticity change synaptic efficacies (Fig. 1D), but in the presence of inhibitory currents, excitatory synapses remain fixed.

### Codependent inhibitory plasticity model

Inhibitory synapses change according to a function *ϕ*_I_(*E, I*; *PRE, POST*) that follows a symmetric STDP curve^13,36^ (Fig. 1E) – synaptic changes are scaled according to the temporal proximity of pre- and postsynaptic spikes. Similar to excitatory plasticity, the learning-rate of inhibitory plasticity is modulated by neighbouring excitatory and inhibitory activity (Fig. 1F,G). In this case, when *E* and *I* currents are equal (*E* = *I*), or when excitatory currents vanish (*E* = 0)^13,14^, there is no change in the efficacy of inhibitory synapses: they remain constant. LTP is induced when excitatory neighbour currents are stronger than inhibitory ones, and vice-versa for LTD. As a consequence, spike times and neighbouring synaptic currents act together but in different timescales, short timescales governed by spikes and long timescale by synaptic currents.

### Stability of excitatory currents

We implemented the above rules in a single leaky integrate-and-fire (LIF) neuron with plastic excitatory synapses that emulate *α*-amino-3-hydroxy-5-methyl-4-isoxazolepropionic acid (AMPA) and NMDA receptors, as well as inhibitory (GABA_A_) synapses (Fig. 2A; see Methods). We initially kept inhibitory synapses fixed and assessed properties of codependent excitatory plasticity alone. We found that our model reproduced the classic experimental results that show the influence of membrane potential depolarisation on synaptic efficacy changes, such that LTD-inducing protocols became LTP-inducing when accompanied by large postsynaptic depolarisation^18^ (Fig. 2B). In the experiment, the switch from LTD to LTP is due to an increase in the magnitude of excitatory currents through NMDA channels for depolarised states. In our model, larger excitatory inputs currents translated into an increased variable *E*, eliciting stronger LTP (see Fig. 1C). Similarly, the interaction of pre- and postsynaptic spikes can also account for efficacy changes based on the frequency of spike pair presentations (Fig. 2C). Notably, in our model, high-frequency of pre- and postsynaptic spike pairs elicited increased LTP (Fig. 2C) due to a direct elevation in excitatory currents (see Fig. 1C). Spike-^6,10^ or voltage-based^7^ models imitate the influence of spike frequency on LTP amplitudes by reacting to an increase in the postsynaptic firing frequency and the consequent increase in spike triplets (post-pre-post). Our model thus varies in the locus of its mechanism: elevated excitatory currents, i.e., a presynaptic driven effect, instead of elevated postsynaptic activity.

**FIG. 2.**
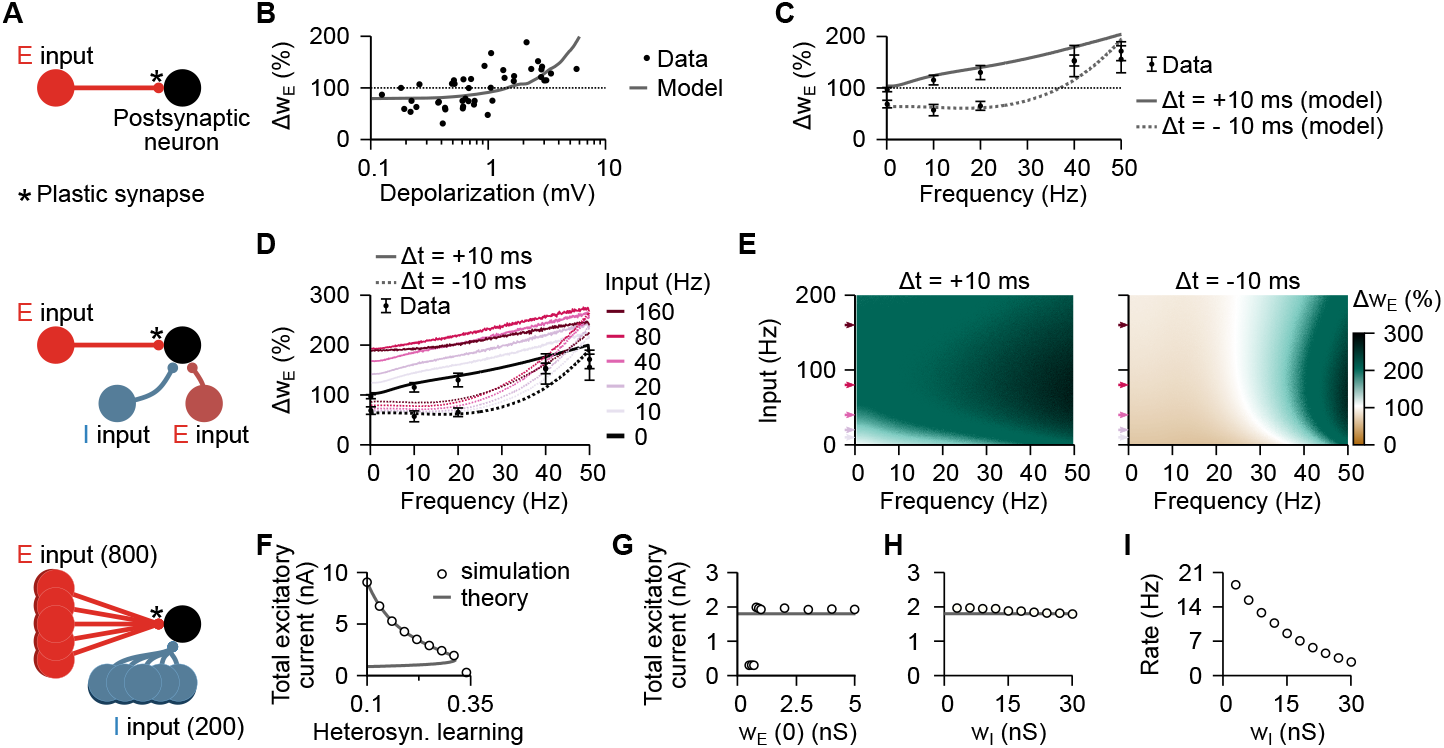
Codependent excitatory synaptic plasticity: influence of voltage and firing-frequency, and excitatory current set-point. **A**, Schematic of the different configurations of the simulation protocols used with codependent excitatory synaptic plasticity. *Top*: Two unidirectly connected excitatory neurons (used in panels B and C). *Middle*: Same as top panel but with extra presynaptic inhibitory and excitatory neurons whose connections are static (used in panels D and E). *Bottom*: A population of 800 excitatory (plastic synapses) and 200 inhibitory (static synapses) neurons connected to a postsynaptic neuron (used in panels F to I). **B**, Changes in excitatory synapses, Δ*w*_E_, as a function of the postsynaptic depolarisation for a 10-ms pre-before-post protocol at 50 Hz. Experimental data points from mouse visual cortex^18^. **C**, Changes in excitatory synapses, Δ*w*_E_, as a function of spike frequency for a spike-timing dependent plasticity protocol with pre-before-post (10 ms) and post-before-pre (−10 ms). Experimental data points from rat visual cortex^17^. Error bars indicate SEM. **D**, Same as panel C for different firing-rates of neighbouring excitatory and inhibitory afferents (colour coded). Plot shows changes in synaptic weight of a single connection while the other two (excitatory and inhibitory) are kept fixed. Excitatory and inhibitory weight of neighbouring synapses were chosen to keep the initial (before plasticity) excitatory and inhibitory currents balanced, and thus same average membrane potential for the same input frequency. **E**, Weight change as a function of input frequency (from neighbouring excitatory and inhibitory synapses; y-axis), and frequency of pairs of spikes (x-axis). Arrows indicate constant input frequencies used in panel D. **F-H**, Total excitatory current (after learning) as a function of the heterosynaptic learning rate (panel F), initial excitatory weights (panel G), and inhibitory weights (panel H). **I**, Firing-rate of the postsynaptic neuron after learning of excitatory synapses for different inhibitory weights for the same simulations in panel H.

In our model, the explicit regulation of plasticity via excitatory and inhibitory currents can alter amplitude, and direction, of synaptic change (Fig. S1A-C). The classic frequency-dependent protocol^17^, for example, has different effects when neighbouring excitatory and inhibitory synapses are simultaneously activated (Fig. 2D,E), highlighting the different roles of neighbouring excitation and inhibition. In contrast, in the traditional spike-^6,10^ or voltage-based^7^ learning rules, neighbouring activation do not affect plasticity as long as it does not influence pre- and postsynaptic spike patterns or the mean postsynaptic membrane potential, i.e., due to balanced excitatory and inhibitory currents (Fig. S2). Moreover, the set-point for the total excitatory current onto a dendritic branch, which emerges from the combination of the Hebbian-LTP and heterosynaptic terms, inherent to our rule (see Fig. 1C, red circle), is determined by the ratio between the learning rate of these two mechanisms (Fig. 2F). The total excitatory current received by a neuron, established by this set-point, is independent of initial excitatory weights and inhibitory input strength (Fig. 2G). With the same amount of excitation – given by the set-point – the more inhibition a neuron receives, the lower its firing-rate (Fig. 2H).

### EI balance and firing-rate set-point

The dynamics of traditional spike-based plasticity rules can be approximated by the firing-rate of pre- and postsynaptic neurons^8,26^. In these types of models, stable postsynaptic activity may be achieved if synaptic weights change towards a firing-rate set-point^8,10^ that controls the dynamics such that excitatory weights increase when the postsynaptic firing-rate is lower than the set-point and decrease otherwise^10,26^. In the same vein, inhibitory weights decrease for low postsynaptic firing-rates (below the set-point) and increase for high firing-rates^8,37^. When both excitatory and inhibitory synapses are plastic (Fig. 3A), the fixed-points from both rules must match to avoid a competition between synapses (Fig. 3B) that would result in synaptic weights to either diverge or vanish (Fig. 3C). Codependent inhibitory plasticity does not have such a problem because there is no firing-rate set-point. Instead, it acts explicitly on excitatory and inhibitory currents (see Fig. 1F,G, and Methods; Fig. S1D), allowing various stable activity regimes for a post-synaptic neuron while avoiding competition with excitatory plasticity (Fig. 3D). A state of global balance between excitation and inhibition, defined by average balance with unspecific correlation between excitation and inhibition for specific input directions (e.g., sound frequency^23^), naturally arises. The complementary spike-based component of codependent inhibitory plasticity (see Fig. 1E) is necessary for the development of a state of detailed balance, defined by correlated excitatory and inhibitory currents for each input direction.

**FIG. 3.**
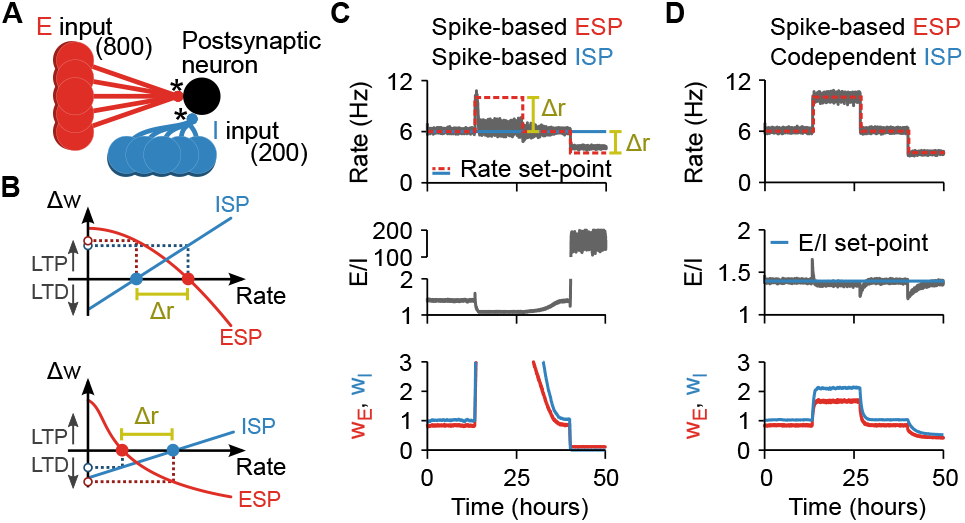
Codependent inhibitory synaptic plasticity: EI balance without firing-rate set-point. **A**, Schematic of the simulations used in panels C and D. A postsynaptic neuron receives 800 excitatory and 200 inhibitory synapses that undergo plasticity, indicate by *. **B**, Schematic of changes in synaptic weight, Δ*w*, as a function of the postsynaptic neuron’s firing-rate for spike-based models with stable set-points. *Top*: Firing-rate set-point from excitatory synaptic plasticity (ESP) is higher than the one from inhibitory synaptic plasticity (ISP). *Bottom*: Firing-rate set-point from ISP is higher than the one from ESP. The interval between the set-point is defined as Δ*r*. **C**, Combination of excitatory^10^ and inhibitory^8^ spike-based rules. *Top*: Firing-rate of a postsynaptic neuron receiving excitatory and inhibitory inputs. Red and blue lines indicate the firing-rate set points imposed by the excitatory^10^ and inhibitory^8^ spike-based learning rules, respectively. The parameters of the learning rules were chosen so that the set-points coincide during the first and third quarters of the simulation. During the second and fourth quarters of the simulation, the set-point imposed by the excitatory spike-based learning rule is increased and decreased, respectively. *Middle*: Ratio between excitatory and inhibitory currents. *Bottom*: Average excitatory (red) and inhibitory (blue) synaptic weights of input neurons normalised to initial value. **D**, Same as panel C for the combination of excitatory spike-based^10^ and codependent inhibitory synaptic learning rules. The blue line in the middle panel indicates the balance set-point imposed by the codependent inhibitory synaptic plasticity.

### Receptive field plasticity

Sensory neurons have been shown to respond more strongly to some features of the stimuli than others, which is thought to facilitate recognition, classification, and discrimination of stimuli. The shape of a neuron’s response profile, i.e., its receptive field, is a result of its input connectivity^23^. Receptive fields are susceptible to change when an animal learns^38^, with strong evidence supporting receptive field changes as a direct consequence of synaptic plasticity^39^.

To assess the functional consequence of codependent plasticity, we studied its performance in receptive field formation for both excitatory and inhibitory synapses jointly. We simulated a postsynaptic LIF neuron receiving inputs from 8 pathways (see Methods) that represent, e.g., different sound frequencies^23^ (Fig. 4A). In this scenario, inhibitory activity acted as a gating mechanism for excitatory plasticity, by keeping the learning rate at a minimum when inhibitory currents were high (see Fig. 1D). Excitatory input weights could thus only change during periods of presynaptic disinhibition, i.e., the learning window (Fig. S3), and were otherwise stable (Fig. 4B,C). In our simulation, we initially set all excitatory weights to the same strength. A receptive field profile emerged at excitatory synapses after a specific sequence of strong stimulation of pathways during the first learning window. The acquired excitatory receptive profile remained stable (static) for hours after the learning period (Fig. 4B, top). Inhibitory synapses changed on a slower timescale (Fig. 4B, bottom) and, due to the spike-timing dependence of codependent ISP, developed a co-tuned field with the excitatory receptive field (Fig. 4D, top). Inspired by experimental work^23^, we then briefly activated a non-preferred pathway during a period of disinhibition (Fig. 4C, top), altering the tuning of excitatory weights and making the previously non-preferred pathway *preferred* (Fig. 4D, middle). This change in tuning happened thanks to the Hebbian component of the codependent excitatory plasticity rule that induced LTP in the active pathway, and the heterosynaptic plasticity component triggering LTD in pathways that were inactive during the learning window. As before, inhibitory weights were reshaped by codependent ISP to a co-tuned field with the most recent excitatory receptive field (Fig. 4C, bottom), reaching a state of detailed balance^3^ (Fig. 4D, bottom). Changes in both excitatory and inhibitory inputs due to codependent plasticity thus reproduced experimental results in rat auditory cortex^23^ (Fig. 4E) without external control.

**FIG. 4.**
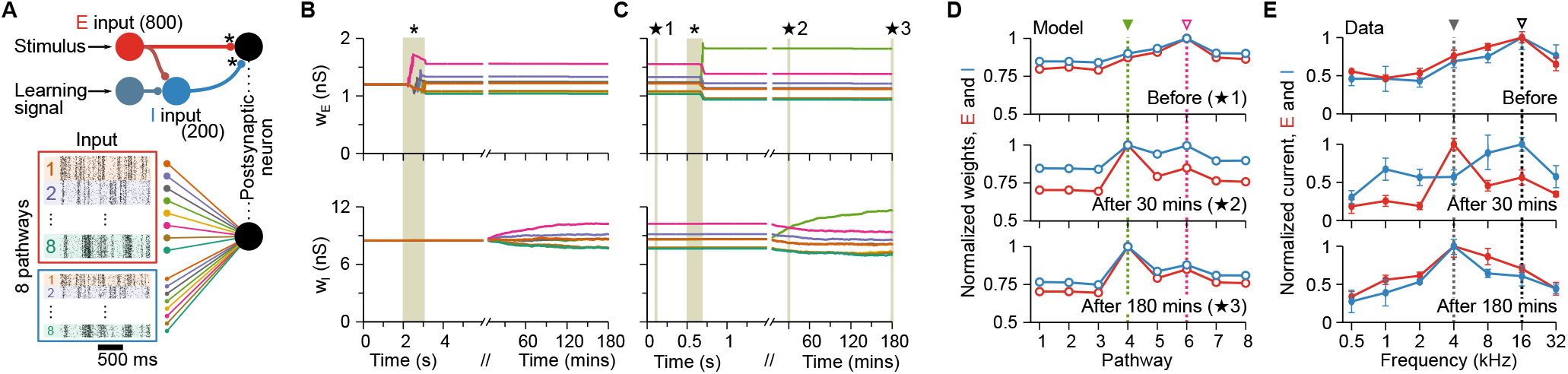
Receptive field plasticity. **A**, External stimulus (e.g., sound) activates a set of correlated excitatory and inhibitory afferents (simulated as inhomogeneous Poisson point processes) that feed forward onto a postsynaptic neuron with plastic synapses, indicated by *. Eight group pathways, consisting of 100 excitatory and 25 inhibitory afferents each, have correlated spike trains. The responsiveness of inhibitory afferents can be modulated by an additional learning signal. **B**, Time course of excitatory (top) and inhibitory (bottom) weights (colour coded by groups). During a “learning window”, indicated by the shaded area and *, all inhibitory afferents are down-regulated. The sequential activation of excitatory input groups (Fig. S3A) in the absence of inhibition establishes a receptive field profile. **C**, Continued simulation from panel B. Weights are stable until inhibition is down-regulated for a 200-millisecond window, indicated by the shaded area and *, during which the green pathway (#4) is activated (Fig. S3B). Consequently, the preferred input pathway switches from #6 (pink) to #4 (green). **D**, Snapshots of the average synaptic weights for the different pathways before (top), immediately after plasticity induction (middle), and at the end of the simulation as indicated by the ★ symbols in panel C. **E**, Experimental data^23^ shows receptive field profiles of excitatory and inhibitory inputs before (top), as well as 30 minutes (middle) and 180 minutes (bottom) after pairing of non-preferred tone and nucleus basalis activation. Error bars indicate SEM.

The formation of stimulus-tuned excitation and co-tuned inhibition was only successful when the learning rules were codependent, and the learning rate of inhibitory plasticity was slow. When excitatory and inhibitory plasticity operated at similar time scales, inhibitory plasticity prevented excitatory weights to change during disinhibition (Fig. 5A, top), because any externally induced decrease in inhibition (disinhibition) was quickly compensated for by inhibitory plasticity (Fig. 5A, bottom). When we reduced the level of codependency, excitatory weights varied wildly, because the modulatory control of inhibitory currents over excitatory plasticity did not effectively decrease the learning rate of excitatory plasticity (Fig. 5B, top). Although a preferred input signal could be momentarily established after the learning window (Fig. 5B, top), the new preference was soon lost because baseline levels of inhibition were not blocking ongoing excitatory plasticity. Inhibitory plasticity quickly compensated the induced disinhibition during the learning period (Fig. 5B, bottom), but inhibitory weights remained unspecific, i.e., without developing a receptive fieldlike profile.

**FIG. 5.**
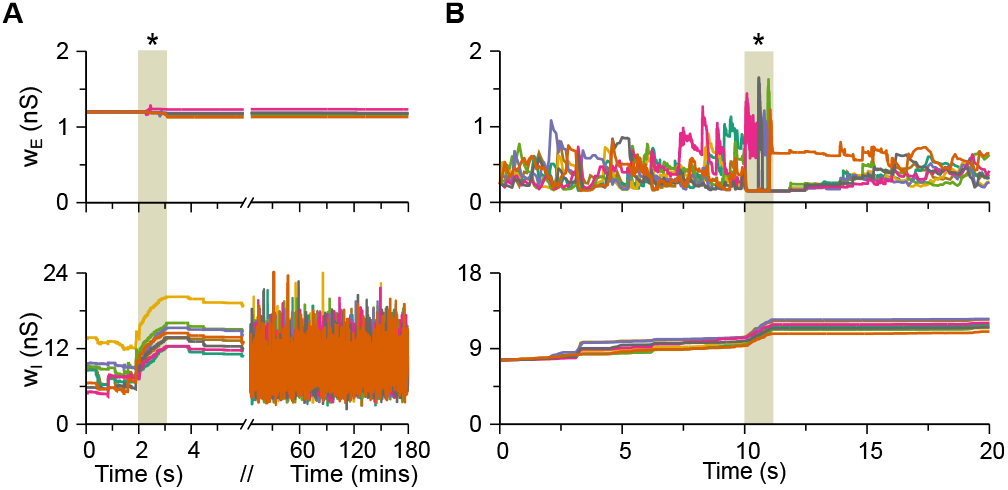
Fast inhibitory plasticity or weak inhibitory control over excitatory plasticity prevents the stable formation of receptive fields. **A**, Same simulation protocol used in Fig. 4B, but with larger learning rate of inhibitory plasticity (increased by 1000-fold). Evolution of excitatory (top) and inhibitory (bottom) weights. The shaded area and * indicate the learning window, when all inhibitory afferents are down-regulated. Excitatory input groups are activated at a specific order for receptive-formation during the learning window (Fig. S3A). **B**, Simulation similar to Fig. 4B, but with weak inhibitory control over excitatory plasticity. The learning window is set between *t* = 10 and *t* = 10.1 seconds with the same levels of disinhibition and excitatory activation sequence as in panel A. Baseline levels of inhibitory currents do not block excitatory plasticity, and thus excitatory weights continuously change.

### Dendritic clustering with single or mixed feature selectivity

The dendritic tree of neurons is a intricate spatial structure that can achieve complex neuronal processing in single neurons that is impossible in single-compartment neuron models^40^. To assess how our learning rules affected the dendritic organisation of synapses, we attached passive dendritic compartments to the soma of our model. Dendritic membrane potentials could be depolarised to values well above the somatic spiking threshold depending on their proximity, i.e., electrotonic distance, to the soma (Fig. 6A). These super-threshold membrane potential fluctuations gave rise to larger NMDA and GABA_A_ current fluctuations in distal dendrites (Fig. 6B). Like in the single compartmental models, when excitation and inhibition were unbalanced (i.e., when receiving uncorrelated inputs), distal dendrites could undergo fast changes due to the current-induced high learning rates for excitatory plasticity (Fig. 6B, thick red line). However, when currents were balanced (i.e., when receiving correlated excitatory and inhibitory inputs), larger inhibitory currents gated excitatory plasticity *off* despite strong excitation, effectively blocking excitatory efficacy changes in these compartments (Fig. 6B, thick blue line). Additionally, the larger distance to the soma, and its consequently weaker passive coupling (Fig. 6C), meant that distal dendrites had a smaller influence on the initiation of postsynaptic spikes.

**FIG. 6.**
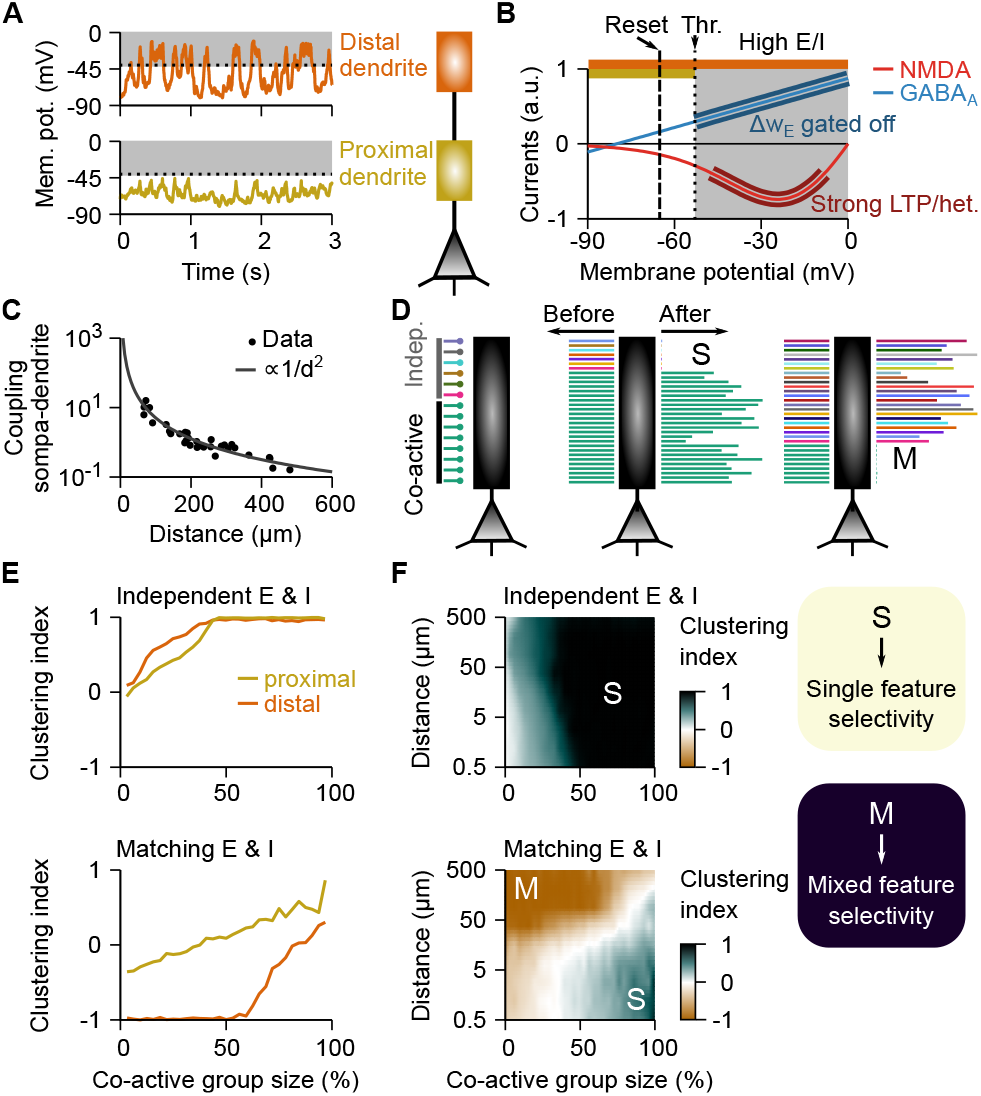
Mixed and single feature selectivity on dendrites depend on presynaptic correlations and distance between soma and dendrite. **A**, Membrane potential fluctuations at distal (top) and proximal (bottom) dendrites during ongoing stimulation. Dashed line shows the spiking threshold at the soma. **B**, NMDA (red) and GABA_A_ (blue) currents as a function of membrane potential. Spiking threshold and reset are indicated by dotted and dashed lines, respectively. **C**, Coupling strength between soma and dendritic branch as a function of electrotonic distance fitted to experimental data^46^. **D**, Schematic of the synaptic organisation onto a dendrite (left). Each line represents a synapse, with co-active synapses bearing the same colour. Examples of clustering of co-active (middle) or independent (right) synapses resulting in single or mixed feature selectivity, respectively, at the level of the dendrite. Line length indicates synaptic weight in arbitrary units. **E**, Clustering index as a function of the size of co-active input group for distal (orange) and proximal (yellow) dendrites with independent (top) and matching (bottom) excitatory and inhibitory inputs. Clustering index is equal to 1 (respectively 1) when only co-active (respectively independent) synapses survived and 0 when all synapses survived (see Methods). **F**, Clustering index (colour-coded) as a function of the size of co-active input group (x-axis) and the distance from the dendrite to the soma (y-axis) for independent (top) and matching (bottom) excitatory and inhibitory inputs. Dark green indicates single feature selectivity while brown indicates mixed feature selectivity.

As a consequence of these two effects – location dependent learning-rates and influence on somatic spike initiation – syn-apses developed differently according to their somatic proximity (Fig. 6D) and according to the activity of their neighbouring inputs. When the majority of excitatory inputs onto a dendritic compartment were co-active, i.e., originated from the same source (representing same stimulus feature such as sound frequency^23^), their co-active synapses were strengthened, creating a cluster of similarly tuned inputs onto a compartment (Fig. 6D, middle). Uncorrelated, independently active excitatory synapses weakened and eventually faded away (Fig. 6D, middle). In contrast, when more than a certain number of excitatory inputs were independent, co-active synapses decreased in weight and faded, while independently active excitatory synapses strengthened (Fig. 6D, right). The number of co-active excitatory synapses necessary for a dendritic compartment to develop single-feature tuning varied with somatic proximity, and whether excitation and inhibition were matched (Fig. S4). Notably, in the balanced state, substantially more coactive excitatory synapses were necessary to create clusters at distal than at proximal dendrites (Fig. 6E), because only large groups of co-active excitatory synapses could initiate LTP-inducing pre-before-post spike pairs. Thus, single-feature selectivity^41,42^ or mixed selectivity^43^ emerged in our model depending on the branch architecture of the dendritic host structure similar to what has been observed experimentally^44,45^ (Fig. 6F).

### Transient amplification in recurrent spiking networks

Up to this point we explored the effects of codependent synaptic plasticity in a single postsynaptic neuron. However, recurrent neuronal circuits typically amplify instabilities of any synaptic plasticity rules at play^10,47^. We thus investigated codependent plasticity in a recurrent neuronal network of 1250 spiking neurons (see Methods; Fig. 7A). In our model, excitatory and inhibitory codependent plasticity allowed the network to self-stabilise in a high-conductance state^48^, with low effective neuronal membrane time-constants (Fig. 7B, left), and strong excitatory connections that were precisely balanced by equally strong inhibition (Fig. 7B, middle, and Fig. S5). Remarkably, the network exhibited a wide distribution of baseline firing-rates (Fig. 7B, right), similar to what has been observed in cortical recordings *in vivo*^49^. To investigate the dynamic behaviour of the network in response to perturbations, we first analysed the learned connectivity matrix of the network with regard to the weights of incoming and outgoing synapses, to identify the neurons which affect the network most strongly (see Methods; Fig. 7C). Next, we delivered step-like stimuli to these neurons, to simulate a large sensory stimulus propagating to other neurons of the network (Fig. 8A,B). We observed several types of responses, ranging from minimal deflections in firing rate to large transient responses, either during the delivery of the stimulus, or in response to its termination. To further quantify these responses, we calculated the average and the *ℓ*_2_-norm of the population (Fig. 8B). Both these measures confirmed that the network was in a regime of ‘transient amplification’^29,50,51^, i.e., a highly responsive state thought to underlie computation-through-population-dynamics^52^ (Fig. 8B). The trajectory of the population dynamics in principle component space showed reliable network wide activity patterns (Fig. 8C) that could be used to control the activity of a readout network with two output units to draw digits (Fig. 8D).

**FIG. 7.**
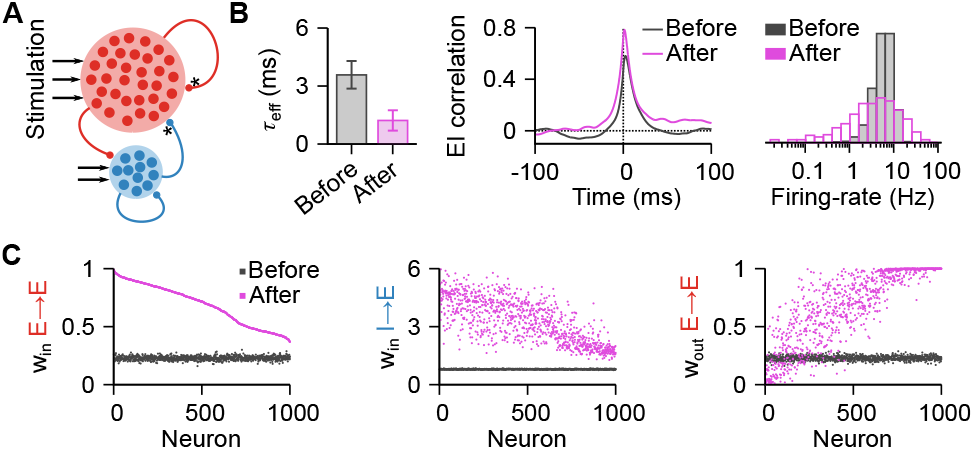
Recurrent network of spiking neurons become highly sensitive after learning period with codependent synaptic plasticity. **A**, Network of 1000 excitatory and 250 inhibitory neurons. Connections between excitatory neurons and from inhibitory to excitatory neurons are plastic (indicated by *). During learning, the external stimulation decreases over time until a state of self-sustained activity is reached. **B**, Summary of changes in excitatory neurons, before (grey) and after (pink) a learning period of 10 hours. *Left*: Effective time constant, calculated as the membrane time constant divided by the sum of all conductances. Error bars indicate SD. *Middle*: Correlation between excitatory and inhibitory currents. *Right*: Firing-rate distribution. **C**, Excitatory and inhibitory connections before (grey) and after (pink) learning. *Left*: Incoming excitatory connections per excitatory neuron. Neurons are ordered from strongest to weakest connection after learning. *Middle*: Incoming inhibitory connections per excitatory neuron. Neurons are ordered as in the left panel. *Right*: Output excitatory connections per excitatory neuron. Neurons are ordered as in the left panel.

**FIG. 8.**
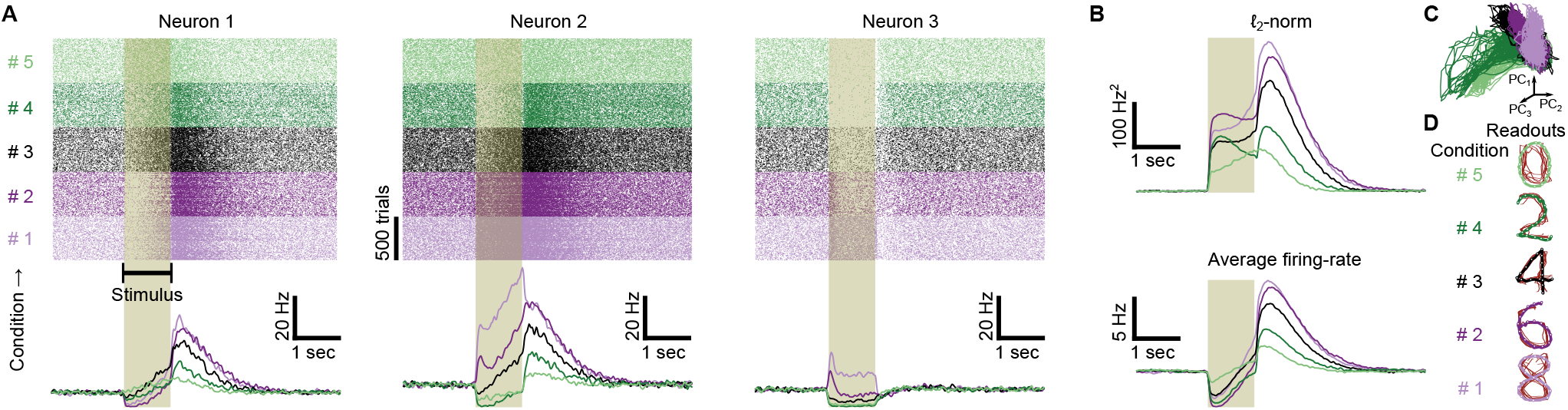
Self-sustained activity and transient amplification in recurrent networks. **A**, Raster plot (top) and mean firing-rate (bottom) of three excitatory neurons for five different stimulation patterns (conditions). **B**, Norm (top) and average firing-rate (bottom) of excitatory neurons for the five stimulation patterns in panel A. **C**, First three principal components of the activity of excitatory neurons from the recurrent network. Each line is the combination of 50 out of 500 trials to compensate for the high trial-to-trial variability. **D**, Output of two readout neurons trained to draw digits on a 2D-plane using the input from the excitatory neurons from the recurrent network (see Methods). Each red thin line is the combination of 50 out of 500 trials (as in panel C) while the thick lines represent the average over the thin lines.

## Discussion

In this paper we introduced a general framework to describe synaptic plasticity as a function of local pre- and postsynaptic interactions, including the modulatory effects of nearby synapses. We built the excitatory and inhibitory plasticity rules according to experimental observations, such that the effect of neighbouring synapses could gate, control, and even invert the direction of efficacy changes^13–20,25^. Importantly, excitatory and inhibitory plasticity were constructed such that they strove towards different fixed points (constant levels of excitatory currents for excitatory plasticity and EI balance for inhibtory plasticity), thus collaborating without mutual antagonism.

In our model, inhibition played an important role in controlling excitatory plasticity, allowing us to make several predictions. First, during periods of disinhibition, inhibitory plasticity has to be slower than excitatory plasticity. A reduction in the amplitude of inhibitory currents directly increased the learning rate of excitatory plasticity, thus allowing for a quick (re-)arrangement of excitatory weights. This effect was prevented if inhibitory weights could immediately increase to compensate the disinhibitory effect, establishing a limit for how fast inhibitory synapses can change. Second, inhibitory control over excitatory plasticity has to be relatively strong. The mechanism that allows excitatory weights to quickly reorganise during periods of disinhibition was also responsible for long-term stability of such modifications when inhibitory activity was at baseline levels. Without strong control, excitatory weights constantly changed due to pre- and postsynaptic activity, drifting away from the synaptic weight pattern established during the period of disinhibition. Finally, our model also predicts that dendrites on which synaptic contacts of both excitatory and inhibitory presynaptic neurons have uncorrelated activity should likely form a connectivity pattern reflecting single-feature selectivity. In this scenario, the initial connectivity pattern will reflect whether a dendritic region may respond to only a few or many input features.

In our model, neighbouring excitatory influence on synaptic plasticity was driven by slow, NMDA-like, excitatory currents, which were required to elicit LTP at excitatory synapses. As a direct consequence, the same pattern of pre- and postsynaptic spike times could produce distinctly different weight dynamics depending on the levels of postsynaptic depolarisation (due to an increase in excitatory currents through NMDA channels caused by the release of the magnesium block^53^). However, an increase in excitatory activity can lead to a rise in the amplitude of excitatory currents (thus also eliciting stronger LTP), even without depolarisation of the postsynaptic neuron (when, e.g., inhibition tightly balances excitation). Postsynaptic membrane potential and presynaptic spike patterns thus independently control the LTP component of excitatory plasticity in our model. This is in line with cooperative view on synaptic plasticity^20^, and experimental findings showing that high-frequency stimulation, which usually elicits LTP, produce LTD when NMDA ion channels are blocked^54^. Further experimental data is necessary to disentangle the specific role of excitatory currents and postsynaptic firing frequency in shaping excitatory synaptic plasticity, and thus unveiling the precise biological form of codependent plasticity.

The set-point dynamics for excitatory currents can be interpreted as a mechanism that normalises excitatory weights by keeping their total combined weights within a range that guarantees a certain level of excitatory currents, similarly to homeostatic regulation of excitatory bouton size in dendrites^55^. Our rule accomplishes this homeostatic regulation through a local combination of Hebbian-LTP and heterosynaptic weakening, similarly to what has been reported in dendrites of visual cortex of mice *in vivo*^21^. Our results show how such plasticity can develop a stable, balanced network that amplifies particular types of input, generating complex spatiotemporal patterns of activity. These networks developed such that they emulate motor-like outputs for both average and single trial experiments^28,56^ without specifically being tuned for it. In our simulations, the phenomenon of transient amplification emerged as a result of the network acquiring a stable high conductance state. This state was established by an autonomous modification of excitatory weights towards a set-point for excitatory currents, balanced by inhibition due to the modification of inhibitory weights towards a regime of precise balance.

Our set of codependent synaptic plasticity rules integrates a number of previously proposed rules that rely on spike times^6,8,10^, synaptic current^9,57^ with implicit voltage dependence^7,58^, heterosynaptic weakening^10^, and neighbouring synaptic activation^32,57^ in a single theoretical framework. In addition to amplifying correlated input activity by way of controlling the efficacy of a synapse, each of the mechanisms in these previous models may replicate a different facet of learning. For example, spike-based plasticity rules can maintain a set of stable firing-rate set-points^8,10,26,27^. Rules based on local membrane potentials^7^, on the other hand, are ideal for spatially extended dendritic structure, making it possible to detect localised activity, and allowing a spatial redistribution of synaptic weights to improve, for example, associative memory when multiple features are learned by a neural network^58^. Similarly, calcium-influx related models^9^ are ideal to incorporate information about presynaptic activation, explaining the emergence of binocular matching in dendrites^57^. Neighbouring activation models^32^ emulate neurotrophic factors that influence the emergence of clustering of synapses during development.

We have unified these disparate approaches in a four-variable model that accounts for the interplay between different synapse types during learning and captures a large range of experimental observations. We have focused only on two types of synapses, i.e., excitatory-to-excitatory and inhibitory-to-excitatory synapses in an abstract setting, but the simplicity of our model allows for the adaptation of a larger number of synaptic types, including, e.g., modulatory signals present in three-factor learning rules^59^. Faithful modelling of a broader range of influences will require additional experimental work to monitor multi-cell interactions by way of, e.g., patterns of excitatory input with glutamate uncaging^60^ or all optical intervention *in vivo*^61,62^. Looking at synaptic plasticity from a holistic viewpoint of integrated synaptic machinery, rather than as a set of disconnected mechanisms, may provide a solid basis to understanding learning and memory.

## Acknowledgements

We thank Chris Currin, Bill Podlaski, and the members of the Vogels group for fruitful discussions. This work was supported by a Research Project Grant by the Leverhulme Trust (RPG-2016-446; EJA), a Sir Henry Dale Fellowship by the Wellcome Trust and the Royal Society (WT100000; EJA, TPV), and a Wellcome Trust Senior Research Fellowship (214316/Z/18/Z; EJA, TPV). For the purpose of open access, the authors have applied a CC BY public copyright license to any authors accepted manuscript version arising from this submission.

## Methods

### Neuron model

#### Point neuron

In the simulations with a postsynaptic neuron described by a single variable (point-neuron) we used a leaky integrate-and-fire (LIF) neuron with after-hyperpolarisation (AHP) current and conductance-based synapses. The postsynaptic neuron’s membrane potential, *u*(*t*), evolved according to a first-order differential equation,

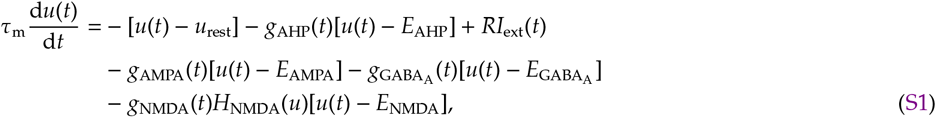

where *τ*_m_ is the membrane time constant (*τ*_m_ = *RC*; leak resistance times membrane capacitance), *u*_rest_ is the resting membrane potential, *g*_AHP_(*t*) is the conductance of the AHP channel with reversal potential *E*_AHP_, *I*_ext_(*t*) is an external current used to mimic experimental protocols to induce excitatory plasticity, and *g*_*X*_(*t*) and *E*_*X*_ are the conductance and the reversal potential of the synaptic channel *X*, respectively, with *X* = {AMPA, NMDA, GABA_A_}. Excitatory NMDA channels were implemented with a nonlinear function of the membrane potential, caused by a Mg^2+^ block, whose effect was simulated via the function

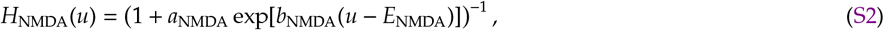

where *a*_NMDA_ and *b*_NMDA_ are parameters. The AHP conductance was modelled as

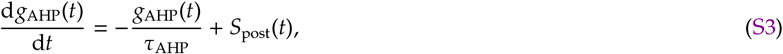

where *τ*_AHP_ is the characteristic time of the AHP channel and *S*_post_(*t*) is the spike train of the postsynaptic neuron,

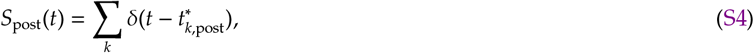

where 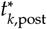 is the time of the k*th* spike of the postsynaptic neuron, and *δ*(·) is the Dirac’s delta. The synaptic conductance was modelled as

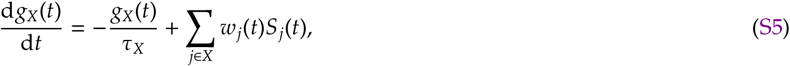

where *τ*_*X*_ is the characteristic time of the neuroreceptor *X*. The sum on the right-hand side of Eq. S5 corresponds to presynaptic spike trains weighted by the synaptic strength *w*_*j*_(*t*). The presynaptic spike train of neuron *j* was modelled as

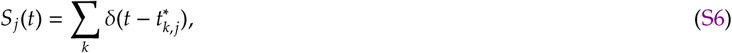

where 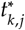 is the time of the k*th* spike of neuron *j*. The postsynaptic neuron elicited an action potential whenever the membrane potential crossed a spiking threshold from below. We simulated two types of threshold: fixed or adaptive.

##### Fixed spiking threshold

A fixed spiking threshold was implemented as a parameter, *u*_th_. When the postsynaptic neuron’s membrane potential crossed *u*_th_ from below, a spike was generated and the postsynaptic neuron’s membrane potential was instantaneously reset to *u*_reset_, and then clamped at this value for the duration of the refractory period, *τ*_ref_. All simulations with a single postsynaptic neuron were implemented with a fixed spiking threshold.

##### Adapting spiking threshold

For the simulations of the recurrent network we used an adapting spiking threshold, *u*_th_(*t*). When the postsynaptic neuron’s membrane potential crossed *u*_th_(*t*) from below, a spike was generated and the postsynaptic neuron’s membrane potential was instantaneously reset to *u*_reset_ without any additional clamping of the membrane potential (the refractory period that results from the adapting threshold is calculated below). Upon spike, the adapting spiking threshold, *u*_th_(*t*), was instantaneously set to 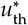, decaying back to its baseline according to

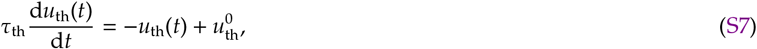

where *τ*_th_ is the decaying time for the threshold variable, and 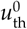 is the baseline for spike generation. The maximum depolarisation of the membrane potential is linked to the reversal potential of NMDA, and thus the absolute refractory period can be calculated as

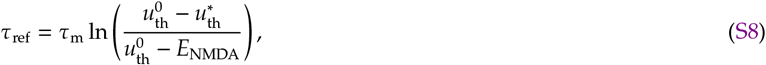

which is the time the adapting threshold takes to decay to the same value as the reversal potential of the NMDA channels. Fig. S6 shows an example of two states that arise from the combination of an adapting threshold and NMDA currents.

#### Two-layer neuron

The two-layer neuron was simulated as a compartmental model with a spiking soma that receives input from *N*_B_ dendritic branches. The soma was modelled as a LIF neuron, and the dendrite as a leaky integrator (without generation of action potentials). Somatic membrane potential evolved according to

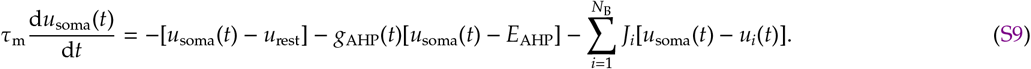

The soma of the two-layer neuron was similar to the point neuron (Eq. S1), however, synaptic currents were injected on the dendritic tree, which interacted with the soma passively through the last term on the right-hand side of Eq. S9, *J*_*i*_ being the conductance that controls the current flow due to connection between the soma and the i*th* dendrite. In Eq. S9, *u*_*i*_(*t*) is the membrane potential of the dendritic branch *i*. When the somatic membrane potential, *u*_soma_(*t*), crossed the threshold, *u*_th_, from below, the postsynaptic neuron generated an action potential, being instantaneously reset to *u*_reset_, and then clamped at this value for the duration of the refractory period, *τ*_ref_.

Dendritic compartments received presynaptic inputs, as well as a sink current from the soma. The membrane potential of the i*th* branch, *u*_*i*_(*t*), evolved according to the following differential equation,

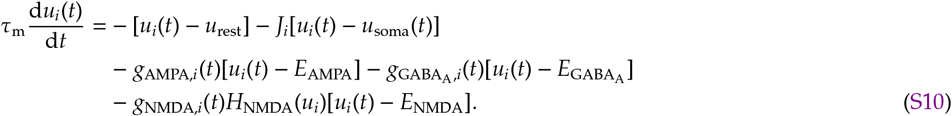

Variables *g*_*X,i*_(*t*), with *X* = {AMPA, NMDA, GABA_A_}, were simulated following Eq. S5. Spikes were not elicited in dendritic compartments, but due to the gating function *H*_NMDA_(*u*) and the absence of spiking threshold, voltage plateaus occurred naturally when multiple inputs arrived simultaneously on a compartment (Fig. 6A). We simulated two compartments with the same coupling with the soma, *J*_*i*_: one whose synapses changed according to the codependent synaptic plasticity model, and one with fixed synapses that acted as a noise source.

##### Coupling strength as function of electrotonic distance

The crucial parameter introduced when including dendritic compartments was the coupling, *J*_*i*_, between soma and the dendritic compartment *i*. Steady changes in membrane potential at the soma are attenuated at dendritic compartments, and this attenuation has been shown to decrease with distance. Without synaptic inputs, and steady membrane potential at both soma and dendritic compartments, Eqs. S9 and S10 are equal to zero, which results in

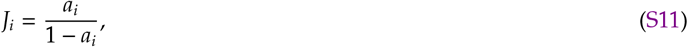

where *a*_*i*_ is the passive dendritic attenuation of the dendritic compartment *i*,

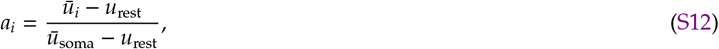

with *ū*_soma_ being a constant steady-state held at the soma, and *ū*_*i*_ being the resulting steady-state at the dendritic compartment *i*. The attenuation is a function of distance as follows,

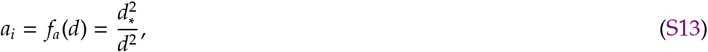

where *d*_*_ is a parameter we fitted from experimental data from Gulledge and Stuart ^46^ (Fig. 6C). We used this fitted parameter in Eq. S11 to approximate the distance to the soma in Fig. 6F according to the soma-dendrite coupling strength used in our simulations.

### Codependent synaptic plasticity model

The codependent plasticity model is a function on both spike times and input currents. We first describe how synaptic currents are accounted and then how excitatory and inhibitory plasticity models were implemented. We defined a variable *E*_*i*_(*t*) to represent the process triggered by excitatory currents at the i*th* dendritic compartment. We considered NMDA currents, which reflect influx of calcium into the postsynaptic cell, as the trigger for biochemical processes that are represented by the state of *E*_*i*_(*t*). Its dynamics is described by

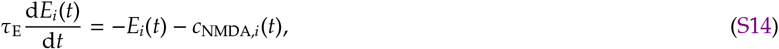

where *τ*_E_ is the characteristic time of the excitatory trace and

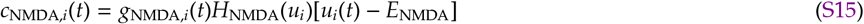

is the excitatory current flowing through NMDA channels. Inhibitory inputs contributed to the plasticity model through a variable *I*_*i*_(*t*). For the inhibitory trace, we used GABA_A_ currents, which reflect influx of chloride, as the trigger of the process described by *I*_*i*_(*t*). The inhibitory trace evolved as

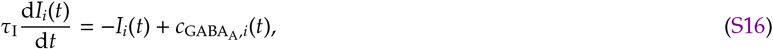

where *τ*_I_ is the characteristic time of the inhibitory trace, 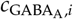 is the inhibitory current described as

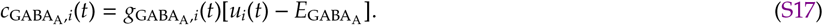

The index *i* is not taken into account for simulations with point-neurons. Notice that both *E*_*i*_(*t*) and *I*_*i*_(*t*) are in units of voltage because the conductance is unit-free in our neuron model implementation (Eq. S1).

### Codependent excitatory synaptic plasticity

The codependent excitatory synaptic plasticity model is a spike-timing-dependent plasticity (STDP) model regulated by excitatory and inhibitory inputs through *E*_*i*_(*t*) and *I*_*i*_(*t*). The weight of the j*th* synapse onto the i*th* dendritic branch, *w*_*ij*_(*t*), changed according to

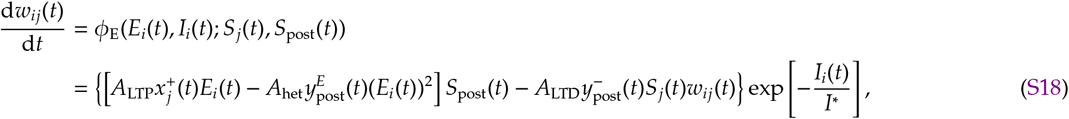

where *A*_LTP_, *A*_het_ and *A*_LTD_ are the learning rates of long-term potentiation, heterosynaptic plasticity and long-term depression, respectively. The additional parameter *I*\* defines the amount of control that inhibitory activity imposes onto excitatory synapses. Variables *S*_post_(*t*) and *S*_*j*_(*t*) represent the postsynaptic and presynaptic spike trains, respectively, as described above for the neuron model (Eqs. S4 and S6). The trace of presynaptic spike train is represented by 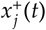, and the traces of postsynaptic spike train (with different time scales) are represented by 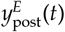 and 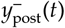. They evolve in time according to

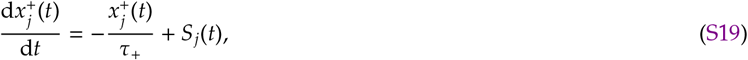

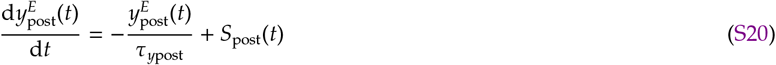

and

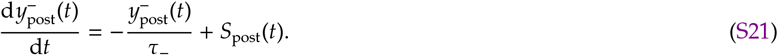

For values of inhibitory trace larger than a threshold, *I*_*i*_(*t*) *> I*_th_, we effectively blocked excitatory plasticity to mimic complete shunting of back-propagating action potentials^35^. We implemented maximum and minimum allowed values for excitatory weights, 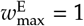 and 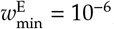, respectively.

### Codependent inhibitory synaptic plasticity

Similar to the excitatory learning rule, the codependent inhibitory synaptic plasticity is a function of spike times and synaptic currents. The weight of the j*th* inhibitory synapse onto the i*th* dendritic compartment, *w*_*ij*_(*t*), changes over time according to a differential equation given by

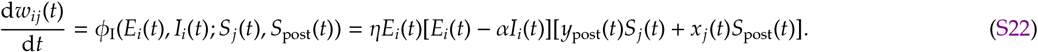

Parameters *η* and *α* control the learning-rate and the balance of excitatory and inhibitory currents, respectively. Variables *y*_post_(*t*) and *x*_*j*_(*t*) are traces of pre and postsynaptic spike-trains, respectively, to create a symmetric STDP-like curve, with dynamics given by

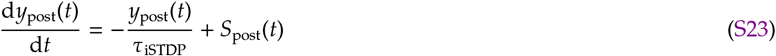

and

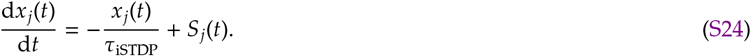

The STDP window is characterised by the time constant *τ*_iSTDP_. We implemented maximum and minimum allowed values for inhibitory weights, 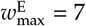 and 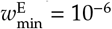, respectively.

### Experimental fit – Fig. 2B,C

We fitted two data sets with the codependent excitatory synaptic plasticity model to asses its dependency on voltage, i.e., membrane potential, and on the frequency of pre- and postsynaptic spikes. We used the same number of pre- and postsynaptic spikes for all our simulations (based on the original experiments). Postsynaptic spikes were induced by the injection of a current pulse, *I*_ext_(*t*) = 3 nA, for the duration of 2 ms.

For Fig. 2B, additional to the current injected for spike generation, we also injected a constant external current to depolarise the neuron’s membrane potential. The protocol consisted in pre- and postsynaptic spikes with 10 millisecond interval, pre-before-post, firing at 50 Hz. The more depolarised the membrane potential, the bigger the effect of the NMDA currents, and therefore more LTP was induced.

The protocol for Fig. 2C consisted on pre- and postsynaptic spikes with either +10 milliseconds (pre-before-post) or −10 milliseconds interval (post-before-pre) in varying firing-rates: 0.1, 10, 20, 40, and 50 Hz. The increase in presynaptic firing-rate caused an increase in NMDA currents which led to strong LTP induction.

Fitting of both protocols was done with brute force parameter sweep on four parameters: *A*_LTP_, *A*_het_, *A*_LTD_, and *τ*_E_.

### Stability

The codependent plasticity model has a rich dynamics that involves changes in synaptic weights due to pre- and postsynaptic spike times, as well as synaptic weight and input currents. In this section we briefly analyse the fixed-points for input currents and synaptic weights for general conditions of inputs and outputs.

Considering separated dendritic compartments we can write the average change in weights (from Eq. S18, ignoring inhibitory inputs) as

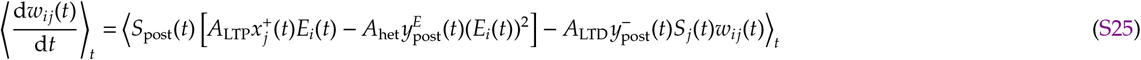

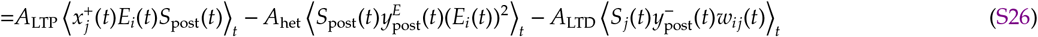

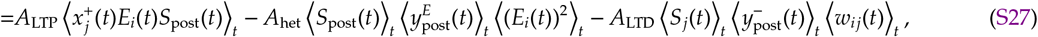

where ⟨·⟩_*t*_ is the average over a long time window, i.e., longer than the timescale of the quantities involved. In Eq. S27 we take into consideration that presynaptic spike times are not influenced by postsynaptic activity, and thus the average of the products in the last term on the right-hand side of Eq. S26 is the equal to the product of the averages. Additionally, we assume no strong correlations between *E*_*i*_(*t*) and *S*_post_(*t*) due to the small fluctuations of the variable *E*_*i*_(*t*). Correlations between pre- and postsynaptic spikes govern the LTP term, and thus cannot be ignored. They also depend on the neuron model, and amount of inhibition a compartment is receiving.

We can conclude from Eq. S27 that the weights from silent presynaptic neurons will vanish due to the heterosynaptic term. In our model, these weights can only vanish in moments of disinhibition, when the inhibitory control over excitatory plasticity is minimum.

For our analysis, we consider that all neurons of the network have nearly stationary firing rates without strong fluctuations. Therefore, the spike trains can be rewritten as average firing rates,

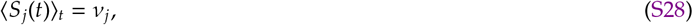

and the traces from the spike trains become

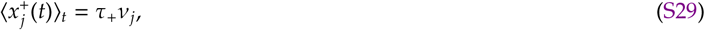

where *ν*_*j*_ is the average firing rate of neuron *j*. The same is valid for the postsynaptic neuron’s firing rate, as well as all other traces.

We first consider the outcome of the excitatory plasticity rule when LTD is not present, *A*_LTD_ = 0, which informs us on steady-state for excitatory currents as a competition between LTP and heterosynaptic plasticity only. In this case, the steady-state of the system is given by

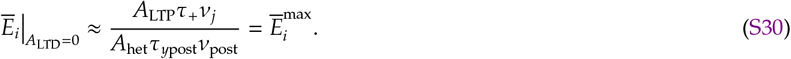

This is also the maximum value for excitatory currents for when LTD is present, as LTD can only decrease synaptic weights. Notice that this fixed-point depends on both pre- and postsynaptic firing-rates, and thus an extra step in necessary to find the fixed-point. For a recurrent network, we can assume that *ν*_*j*_ = *ν*_post_ and thus

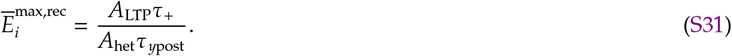

Notice that the maximum excitatory current onto a neuron embedded in a recurrent network is independent on firing-rate of pre- and postsynaptic neurons.

To have an idea of the contribution of LTD for the excitatory inputs, we used a variant of the model in which LTD does not depend on the weight, *w*_*ij*_(*t*), and heterosynaptic plasticity does not depend on the postsynaptic trace, 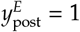. We thus find that

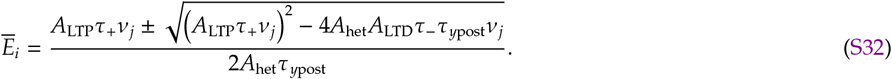

In Fig. 2F we simulated this version of the model with constant presynaptic firing-rate, *ν*_*j*_, and plotted the theory as Eq. S32. To test the effect of input firing-rate and LTD with weight dependency, we simulated the protocol as in Fig. 2F, but with different levels of excitatory input, LTD, and inhibitory gating (Fig. S7). These simulations (Fig. S7) show that, although the input has an effect on the fixed-point of excitatory currents, this effect is minimal compared to the effect of the learning-rates.

Applying the same idea to the codependent inhibitory synaptic plasticity model, we get the following average dynamics for the inhibitory weight connecting presynaptic neuron *j* and the dendritic compartment *i*,

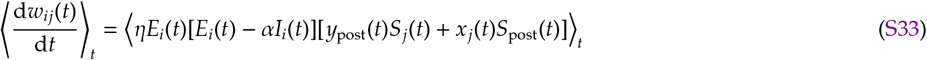

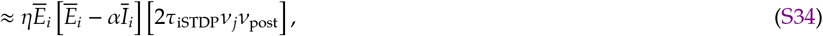

where 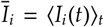. From Eq. S34, we get the steady-state for the inhibitory learning rule, which results in the balance between excitation and inhibition given by *α*,

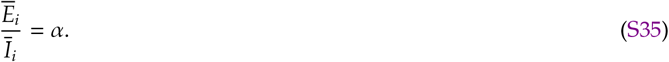

### Synaptic changes for simple spike patterns and fixed excitatory/inhibitory input levels

From Eq. S18 and Eq. S22 we calculated changes in excitatory and inhibitory synapses for simple spike patterns (Fig. S1). We considered fixed excitatory and inhibitory inputs, *E* and *I*, respectively, and calculated changes in a given excitatory synapse as

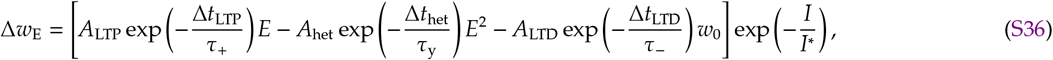

where Δ*t*_LTP_ is the interval between pre- and postsynaptic spikes (pre-before-post), Δ*t*_het_ is the interval between two consecutive postsynaptic spikes, and Δ*t*_LTD_ is the interval between post- and presynaptic spikes (post-before-pre). In a similar fashion, we calculated changes at a given inhibitory synapse as

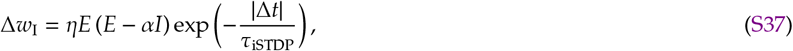

where Δ*t* is the interval between pre- and postsynaptic spikes, being positive for pre-before-post and negative for post-before-pre spike patterns.

### Inputs

#### Single output neuron (feedforward network)

Presynaptic spike trains for single neurons were implemented as follows. A spike of neuron *j* occurs in a given time-step of duration Δ*t* with probability *p*_*j*_(*t*) if there was no spike elicited during the refractory period beforehand, 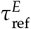 for excitatory and 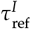 for inhibitory inputs, respectively, and zero otherwise. For a constant probability *p*_*j*_(*t*) = *p*_*j*_ the mean firing-rate, *ν*_*j*_, is therefore

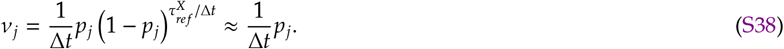

Different simulation paradigms (or tests) are defined by the input statistics, which are described below. When pathways are included, the probability of having a presynaptic spike changes over time to create correlations between different presynaptic inputs.

##### Constant firing-rate – Fig. 2F-I, Fig. 3, and Fig. S7

When constant background firing-rate was simulated, presynaptic neurons fired action potentials with probability *p*_*j*_ = 5 × 10^−5^ for excitatory and *p*_*j*_ = 10^−4^ for inhibitory neurons, resulting in *ν*_*j*_ ≈ 4.88 Hz for excitatory and *ν*_*j*_ ≈ 9.51 Hz for inhibitory neurons, considering that simulations were implemented with a time step Δ*t* = 0.1 ms and that refractory periods were 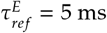 and 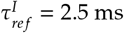.

##### Constant firing-rate – Fig. 2D,E and Fig. S2

To explore how neighbouring neurons would affect plasticity at a given synapse we simulated the same protocol from the frequency-dependency experiment^17^ (Fig. 2C) with the addition of one excitatory and one inhibitory presynaptic neuron active with different firing-rates (Fig. 2D,E). These extra synapses were kept fixed during the simulation and simulated without refractory period 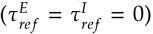, and we tested different firing-rates (from 0 to 200 Hz). We simulated the codependent excitatory plasticity rule with (Fig. 2D,E and Fig. S2A) and without (Fig. S2B) inhibitory gating, and compared with spike-based^6,10^ (Fig. S2C) and voltage-based^7^ (Fig. S2D) learning rules (described below).

##### Variable firing-rate (pathways) for receptive-field plasticity – Fig. 4, Fig. 5, and Fig. S3

For correlated inputs (defined as pathways), we generated spike trains using an inhomogeneous Poisson process. A pathway is defined as a group of 100 excitatory and 25 inhibitory afferents (spike trains of presynaptic neurons) with two components: a constant background firing-rate and a fluctuating firing-rate taken from an Ornstein-Uhlenbeck (OU) process as described below. The background firing-rate for all 800 excitatory and 200 inhibitory afferents was given by a probability of 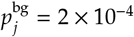 for excitatory and 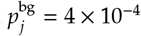 for inhibitory afferents, with respective background firing-rates of 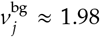 and 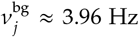 for excitatory and inhibitory presynaptic neurons, respectively, considering a time step Δ*t* = 0.1 ms and refractory periods of 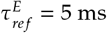 and 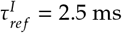.

The fluctuating firing-rate of the pathway *μ* was created from an OU process. We used an auxiliary variable, *y*_*μ*_(*t*), that followed stochastic dynamics given by

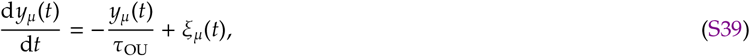

where *τ*_OU_ is the time-constant of the OU process, and *ξ*_*μ*_(*t*) is a random variable drawn from a Gaussian distribution with zero-mean and unitary standard deviation. The fluctuating firing-rate was then defined as

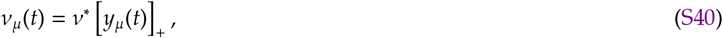

where *ν*^*^ = 250 Hz is the amplitude of the fluctuations and [·]_+_ is a rectifying function. The probability of a spike generated because of the fluctuating firing-rate is thus 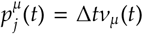. The probability of a presynaptic afferent *j* belonging to pathway *μ* due to both background and fluctuating firing-rate was given by

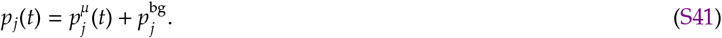

In Fig. 4 we have two learning windows: first to learn the initial receptive field profile (Fig. 4B), and later to learn the new configuration of the receptive field profile (Fig. 4C). During both learning periods, we set the firing-rate of all inhibitory neurons to 40% of background firing-rate, and the inactive excitatory pathways to background firing-rate. The first learning period lasts 1.1 seconds and has the following sequence of activation of excitatory groups: 0.5 seconds for pathway number 6, followed by 0.2 seconds of pathway number 7, 0.2 seconds of pathway number 5, 0.1 seconds of pathway number 4, and finally 0.1 seconds of pathway number 8 (Fig. S3A). The second learning period lasts 0.2 seconds and we only activated pathway number 4 (Fig. S3B). Activation of a pathway consisted in increasing the firing-rate probability to *p*_*j*_ = 0.01, resulting in a firing-rate of *ν*_*μ*_ ≈ 60 Hz.

##### Variable firing-rate (pathways) for dendritic clustering – Fig. 6 and Fig. S4

To explore the clustering effect on dendritic compartments we divided the input spikes in pathways to have co-active or independent presynaptic afferents. We used the same implementation as for the *Variable firing-rate (pathways) for receptive field plasticity* above, but we changed the number of afferents per group in both excitatory and inhibitory presynaptic inputs. A dendritic compartment received 32 excitatory and 16 inhibitory afferents. In Fig. 6E, Fig. 6F, and Fig. S4 we used two conditions: independent E & I, and matching E & I. In both cases, the number of excitatory afferents following the same fluctuating firing-rate was increased from 1 (0% co-active group size) to 32 (100% co-active group size), while the remaining excitatory afferents had independent fluctuating firing-rates. For independent excitatory and inhibitory inputs (independent E & I), all 16 inhibitory afferents followed independent fluctuating firing-rates. For matching excitatory and inhibitory inputs (matching E & I), 8 inhibitory afferents followed the same fluctuations in firing-rate as the co-active excitatory group (of different sizes), while the other 8 inhibitory afferents were independent.

#### Recurrent network

The simulation with the recurrent network had two parts, the learning period with both excitatory and inhibitory plasticity active, and the recall period without plasticity mechanisms active.

##### Learning period

During the beginning of the learning period of *T* = 10 hours (Fig. S5) we kept the network receiving a minimum of external input to avoid inactivity. The implementation of presynaptic spike train was as follows. In the beginning of the simulation (first 1 hour of simulated time), each excitatory neuron of the network received a spike train from one external source with constant probability *p* = 0.01 (time step Δ*t* = 0.1 ms) to mimic 100 presynaptic afferents firing at 1 Hz. We decreased the probability *p* = 0.001 for another 1 hour of simulated time, and then set it to zero. External inputs only affected AMPA conductances to keep the external influence on the network’s dynamics minimal.

##### Recall period

To elicit transient amplification, we selected specific neurons to receive external input based on the resulted weight matrix and the neurons’ baseline firing-rate. Before and after stimulation, no external input was implemented, meaning that the network was in a state of self-sustained activity. During the stimulation period, network neurons were stimulated with presynaptic spikes with a constant firing-rate with different amplitudes for each of the 5 conditions shown in Fig 8. We ordered excitatory and inhibitory neurons according to their baseline firing-rate multiplied by total output weight (from maximum to minimum values), 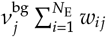 for excitatory and 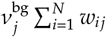 for inhibitory neurons, where *N*_E_ is the total number of excitatory neurons, and *N* is the total number of neurons in the recurrent network. We assumed that the bigger the baseline firing-rate multiplied by the output weight, the bigger the neuron’s influence on the rest of the network.

Considering the the order of maximal (max) to minimal (min) influence we used the following patterns of stimulation. For condition #1, external firing-rate was decreased from 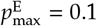 to 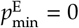 for excitatory neurons, and followed the opposite order for inhibitory neurons. For condition #2, external firing-rate was increased from 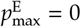 to 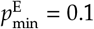 for excitatory neurons, and followed the same order for inhibitory neurons. For condition #3, external firing-rate was decreased from 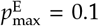 to 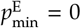 for excitatory neurons, and followed the same order for inhibitory neurons. For condition #4, external firing-rate was increased from 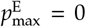 to 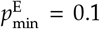 for excitatory neurons, and followed the opposite order for inhibitory neurons. For condition #5, external firing-rate was chosen randomly from a uniform distribution between 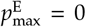 and 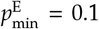 for excitatory neurons, and similarly for inhibitory neurons. Notice that the pattern of stimulation that activated excitatory and inactivated inhibitory neurons with large impact on the network (condition #1), amplification was the largest amongst the conditions, and when the pattern of stimulation was random (condition #5) the resulting network dynamics had minimum amplification (Fig. 8A,B).

### Clustering index for dendritic dynamics - Fig. 6E,F

We defined the clustering index as

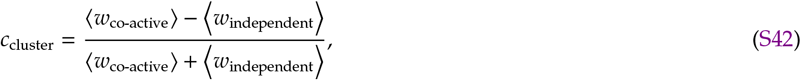

where ⟨*w*_co-active_⟩ is the average of the weights from the co-active excitatory group, and ⟨*w*_independent_⟩ is the average of the weights from all independent groups after learning (see individual weight dynamics in Fig. S4). When *c*_cluster_ = 1 the excitatory weights from the co-active group survived after learning, and independent ones vanished, while for *c*_cluster_ = −1 the opposite happened. Both co-active and independent groups survived after learning when *c*_cluster_ ≈ 0.

### Training an output to draw digits

We connected all excitatory neurons of our recurrent network to a two-layer network of nonlinear units. The second layer was connected to two nonlinear readouts that represented movement in the horizontal *x*(*t*) and vertical *y*(*t*) directions of a 2D plane. The dynamics of the i*th* unit of the first layer (total number of units *N*_1_ = 50) was simulated as

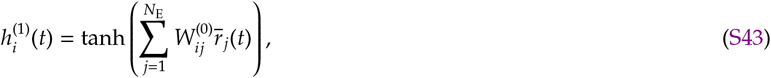

where *W*_*ij*_ is the connection between neuron *j* from the network and the i*th* unit from the first layer and 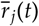 is the average firing-rate of neuron *j* (over 50 trials, see below). The dynamics of neuron *j* for an individual trial is given by

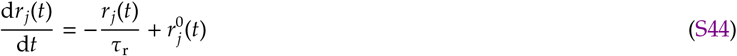

with

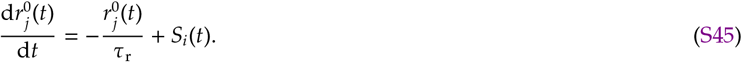

The firing-rate, *r*_*i*_(*t*), is the spike train of neuron *i* (Eq. S6) filtered twice (double exponential filter) with the time constant *τ*_r_ = 10 ms. As input to the first layer we used 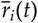 which is the average of *r*_*i*_(*t*) over 50 random trials out of 500 that were simulated. The second layer received input from the first layer, and the dynamics of the i*th* unit (total number of units *N*_2_ = 25) of the second layer was simulated as

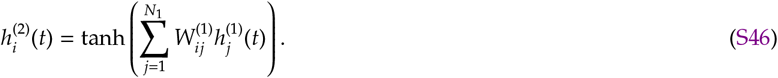

The two readouts were simulated as

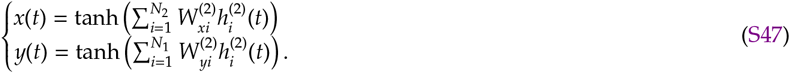

The time-course of the simulation was divided into 35 bins of equal interval (Δ*T* = 30 ms), and the period after the stimulus onset was used to train the output weights to draw five digits: 0, 2, 4, 6, and 8, for the five different conditions from Fig. 7 that resulted from distinct patterns of stimulation. Each training epoch (single digit presentation) was simulated with the average firing-rate of 50 trials, randomly selected out of 500 to account for the large trial-to-trial variability of the spike patterns of each trial, mimicking a larger network in which groups of 50 neurons have a similar activity pattern. We used the same activity patterns fed to the two-layer network, 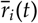, to compute the principal components shown in Fig. 8C. All the weights were trained using gradient descent to minimise the square error between output and the drawing. Fig. 8D shows the average 2D trajectory for each digit in thick line, and 10 individual epochs in red lines. We did not perform any benchmark test as this is beyond the scope of this paper.

### Comparison to other learning rules

We used spike-based learning rules to explore the balancing mechanisms of the codependent inhibitory synaptic plasticity rule (Fig. 3). Additionally, we compared the excitatory codependent synaptic plasticity model with a spike- and a voltage-based learning rule when extra excitatory and inhibitory inputs are received by the postsynaptic neuron (Fig. S2).

### Spike-based plasticity rules

We implemented spike-based learning rules that have been developed to impose a set-point for postsynaptic neurons. In Fig. 3C, we combined excitatory and inhibitory spike-based plasticity rules to show how they can destructively compete when their firing-rate set-points do not match. In Fig. 3D, we combined an excitatory spike-based plasticity rule with the codependent inhibitory synaptic plasticity rule to show how the competition is not present when the plasticity rules dynamics follow fixed-points for different quantities – here ESP imposes a firing-rate set-point while ISP imposes an input currents set-point. We also compared the spike-based excitatory plasticity rule to the codependent excitatory plasticity rule for the same protocol in Fig. 2C, but adding activity of an excitatory and an inhibitory neuron with different firing-rates (Fig. S2C).

#### Spike-based excitatory plasticity rule

The excitatory plasticity rule was based on Pfister and Gerstner ^6^ and Zenke *et al*. ^10^. We used a modified version from Zenke *et al*. ^10^ to create a stable-fixed point for the postsynaptic firing-rate. The weight from neuron *j* to neuron *i* changes according to

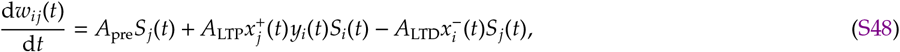

where *A*_pre_, *A*_LTP_, and *A*_LTD_, are the learning rates for the neurotransmitter induced plasticity, long-term potentiation, and long-term depression, respectively. The spike train of pre- and postsynaptic neurons, *S*_*j*_(*t*) and *S*_*i*_(*t*), respectively, are defined by Eq. S6 and Eq. S4. The traces 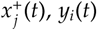, *y*_*i*_(*t*), and 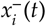 followed the same dynamics described in Eq. S19, with time constants *τ*_+_, *τ*_*y*_ and *τ*_−_ respectively. The learning rule described by Eq. S48 has a stable fixed-point for low postsynaptic firing-rate. The stable firing-rate set point was given by

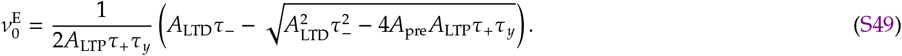

To change the firing-rate set-point during the simulation in Fig. 1F, we changed *A*_pre_, keeping all the other parameters fixed.

#### Spike-based inhibitory plasticity rule

The spike-based inhibitory plasticity rule was based on Vogels *et al*. ^8^. This inhibitory plasticity rule follows a Hebbian-like shape that imposes a postsynaptic firing-rate set-point by balancing excitation and inhibition. An inhibitory weight from neuron *j* to neuron *i* evolved as

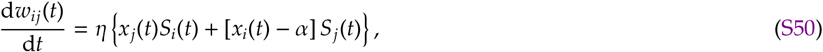

where *η* is the learning rate, *x*_*j*_(*t*) and *x*_*i*_(*t*) are traces of pre- and postsynaptic spike trains, respectively, and they evolve with the same time constant *τ*_iSTDP_. This learning rule is known to result in a stable firing-rate set-point for postsynaptic neurons given by

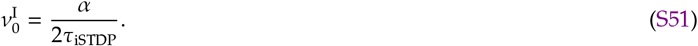

#### Voltage-based excitatory plasticity rule

We also compared the codependent excitatory synaptic plasticity model with a voltage-based model for a frequency dependent protocol^17^ when additional presynaptic activity is included (Fig. S2D). The excitatory weight of the connection from neuron *j* to neuron *i* changed according to

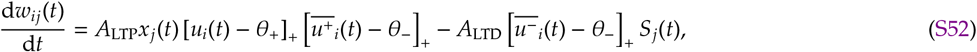

where *A*_LTP_ and *A*_LTD_ are the learning rates for LTP and LTD, respectively, *x*_*j*_(*t*) is the trace of the presynaptic spike train, *θ*_+_ is the voltage threshold for LTP, *θ*_−_ is the voltage threshold (for the voltage low pass-filter) for LTP and LTD. The trace of the presynaptic spike train is given by

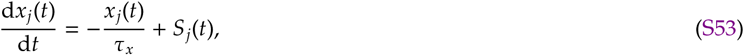

where *τ*_*x*_ is the time constant for the trace. The voltage-based learning rule depends on the low-pass filter of the membrane potential, which creates a dependency on the recent history of membrane potential. There are two low-pass filters in the voltage-based model: one for LTP, 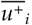, and one for LTD, 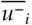, with dynamics given by

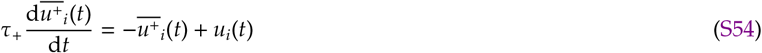

and

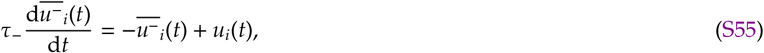

where *τ*_+_ and *τ*_−_ are the time constants for the voltage low-pass filters for LTP and LTD, respectively.

To account for the plasticity inducing effects of the action potential, we modified the LIF neuron model so that it had a short depolarised period corresponding to the action potential. Following the implementation from the subsection *Point neuron - Fixed spiking threshold*, when the membrane potential crossed the threshold, *u*_th_ from below, the membrane potential was instantaneously set to *u*_*AP*_ = 30 mV, and it was then clamped at this voltage for the duration of the action potential *τ*_AP_ = 2 ms. After the duration of the action potential, the membrane potential was instantaneously reset to 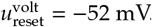.

### Simulations

All simulations were run with Intel Fortran. Parameters used in simulations are defined in Tables S1, S2, S3, S4, S5, S6, S7, S8, and S9.

## Parameters

**TABLE S1.**
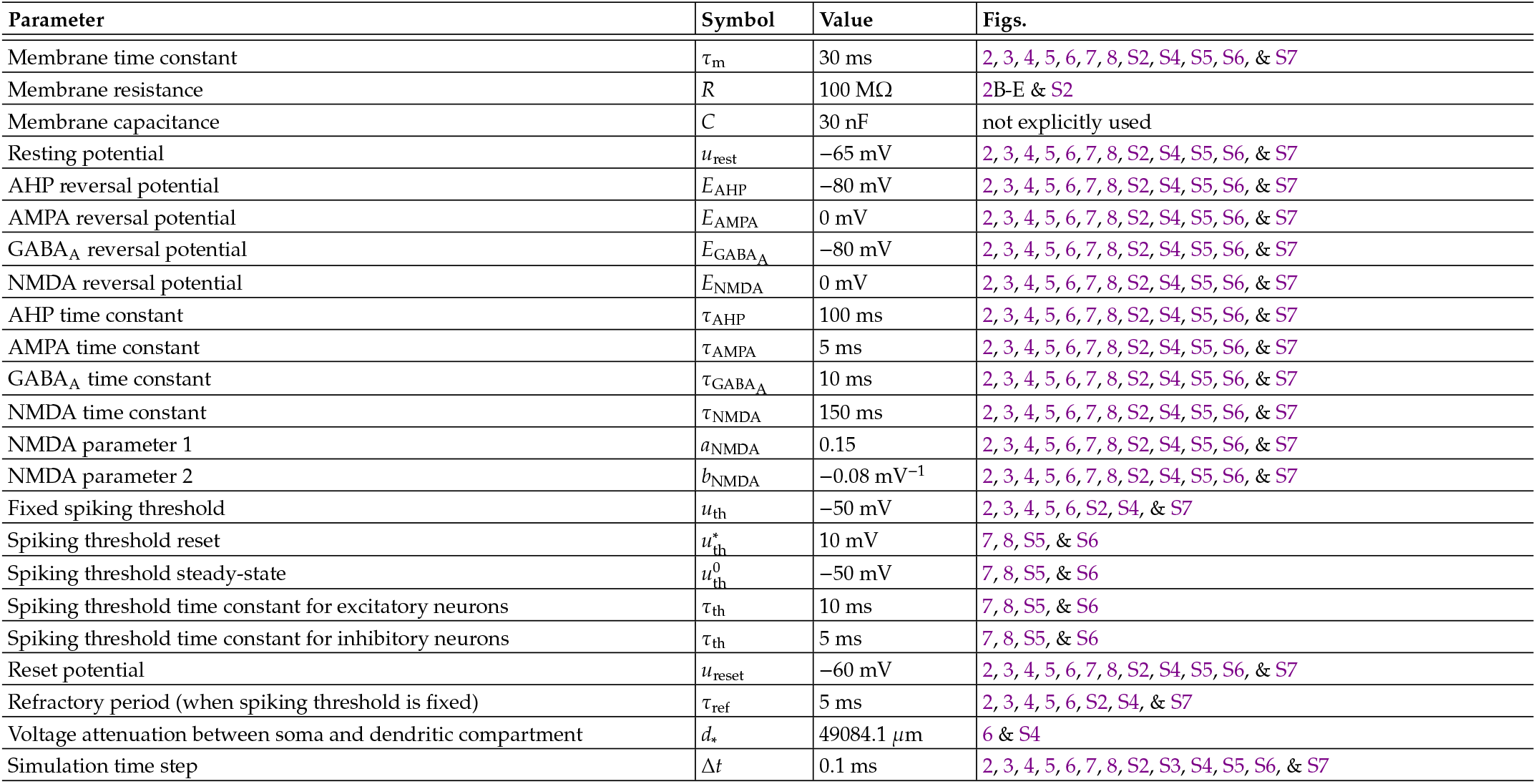
Simulation parameters for the leaky integrate-and-fire neuron.

**TABLE S2.**
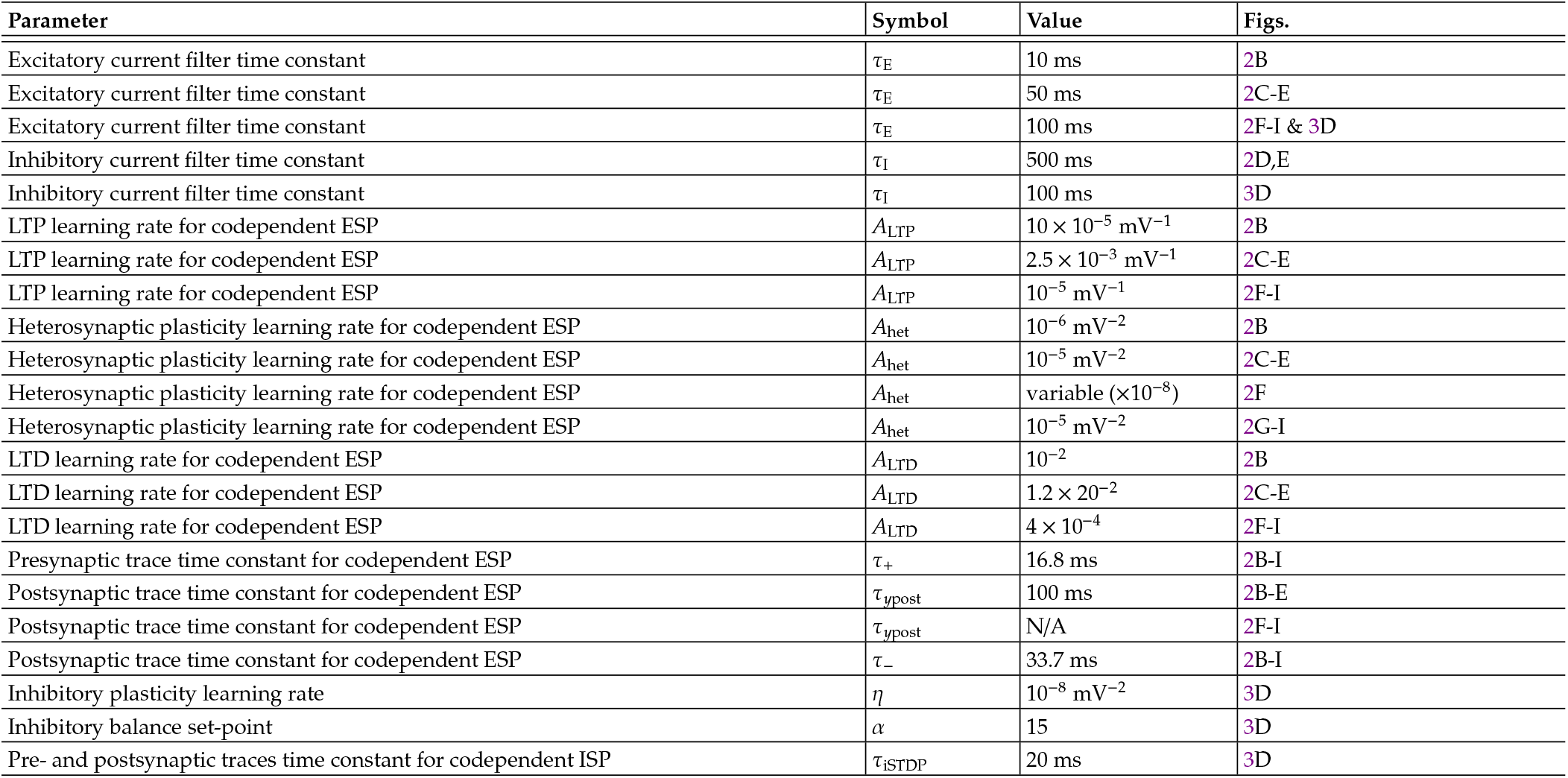
Simulation parameters for codependent synaptic plasticity model in Fig. 2 and Fig. 3.

| Parameter | Symbol | Value | Figs. |
| --- | --- | --- | --- |
| Excitatory current filter time constant | $\tau_E$ | 10 ms | 2B |
| Excitatory current filter time constant | $\tau_E$ | 50 ms | 2C-E |
| Excitatory current filter time constant | $\tau_E$ | 100 ms | 2F-I & 3D |
| Inhibitory current filter time constant | $\tau_I$ | 500 ms | 2D,E |
| Inhibitory current filter time constant | $\tau_I$ | 100 ms | 3D |
| LTP learning rate for codependent ESP | $A_{LTP}$ | $10 \times 10^{-5}$ mV<sup>-1</sup> | 2B |
| LTP learning rate for codependent ESP | $A_{LTP}$ | $2.5 \times 10^{-3}$ mV<sup>-1</sup> | 2C-E |
| LTP learning rate for codependent ESP | $A_{LTP}$ | $10^{-5}$ mV<sup>-1</sup> | 2F-I |
| Heterosynaptic plasticity learning rate for codependent ESP | $A_{het}$ | $10^{-6}$ mV<sup>-2</sup> | 2B |
| Heterosynaptic plasticity learning rate for codependent ESP | $A_{het}$ | $10^{-5}$ mV<sup>-2</sup> | 2C-E |
| Heterosynaptic plasticity learning rate for codependent ESP | $A_{het}$ | variable ($\times 10^{-8}$) | 2F |
| Heterosynaptic plasticity learning rate for codependent ESP | $A_{het}$ | $10^{-5}$ mV<sup>-2</sup> | 2G-I |
| LTD learning rate for codependent ESP | $A_{LTD}$ | $10^{-2}$ | 2B |
| LTD learning rate for codependent ESP | $A_{LTD}$ | $1.2 \times 10^{-2}$ | 2C-E |
| LTD learning rate for codependent ESP | $A_{LTD}$ | $4 \times 10^{-4}$ | 2F-I |
| Presynaptic trace time constant for codependent ESP | $\tau_+$ | 16.8 ms | 2B-I |
| Postsynaptic trace time constant for codependent ESP | $\tau_{ypost}$ | 100 ms | 2B-E |
| Postsynaptic trace time constant for codependent ESP | $\tau_{ypost}$ | N/A | 2F-I |
| Postsynaptic trace time constant for codependent ESP | $\tau_-$ | 33.7 ms | 2B-I |
| Inhibitory plasticity learning rate | $\eta$ | $10^{-8}$ mV<sup>-2</sup> | 3D |
| Inhibitory balance set-point | $\alpha$ | 15 | 3D |
| Pre- and postsynaptic traces time constant for codependent ISP | $\tau_{ISTDP}$ | 20 ms | 3D |

**TABLE S3.** Simulation parameters for spike-based synaptic plasticity models in Fig. 3.

| Parameter | Symbol | Value | Figs. |
| --- | --- | --- | --- |
| Neurotransmitter induced plasticity learning rate for spike-based ESP | $A_{pre}$ | $[9.4, 15, 9.4, 5] \times 10^{-3}$ | 3C,D |
| LTP learning rate for spike-based ESP | $A_{LTP}$ | $10^{-3}$ | 3C,D |
| LTD learning rate for spike-based ESP | $A_{LTD}$ | $4.5 \times 10^{-2}$ | 3C,D |
| Presynaptic trace time constant for spike-based ESP | $\tau_+$ | 16.8 ms | 3C,D |
| Postsynaptic trace time constant for spike-based ESP | $\tau_y$ | 100 ms | 3C,D |
| Postsynaptic trace time constant for spike-based ESP | $\tau_-$ | 33.7 ms | 3C,D |
| Spike-based inhibitory plasticity learning rate | $\eta$ | $10^{-3}$ | 3C |
| Spike-based inhibitory plasticity LTD | $\alpha$ | 0.228 | 3C |
| Pre- and postsynaptic traces time constant for codependent ISP | $\tau_{iSTDP}$ | 20 ms | 3C |

**TABLE S4.** Simulation parameters for codependent synaptic plasticity model in Fig. 4 and Fig. 5.

| Parameter | Symbol | Value | Figs. |
| --- | --- | --- | --- |
| Excitatory current filter time constant | $\tau_E$ | 10 ms | 4 & 5 |
| Inhibitory current filter time constant | $\tau_I$ | 100 ms | 4 & 5 |
| LTP learning rate for codependent ESP | $A_{LTP}$ | $5 \times 10^{-4} \text{ mV}^{-1}$ | 4 & 5 |
| Heterosynaptic plasticity learning rate for codependent ESP | $A_{het}$ | $10^{-8} \text{ mV}^{-2}$ | 4 & 5 |
| LTD learning rate for codependent ESP | $A_{LTD}$ | 0.5 | 4 & 5 |
| Inhibitory gating term for codependent ESP | $I^*$ | 60 mV | 4 |
| Inhibitory gating term for codependent ESP | $I^*$ | 600 mV | 5 |
| Inhibitory threshold for codependent ESP | $I_{th}$ | 200 mV | 4 |
| Inhibitory threshold for codependent ESP | $I_{th}$ | 2000 mV | 5 |
| Presynaptic trace time constant for codependent ESP | $\tau_+$ | 16.8 ms | 4 & 5 |
| Postsynaptic trace time constant for codependent ESP | $\tau_{ypost}$ | 100 ms | 4 & 5 |
| Postsynaptic trace time constant for codependent ESP | $\tau_-$ | 33.7 ms | 4 & 5 |
| Inhibitory plasticity learning rate | $\eta$ | $10^{-9} \text{ mV}^{-2}$ | 4 5 |
| Inhibitory balance set-point | $\alpha$ | 0.94 | 4 & 5 |
| Pre- and postsynaptic traces time constant for codependent ISP | $\tau_{iSTDP}$ | 20 ms | 4 & 5 |

**TABLE S5.**
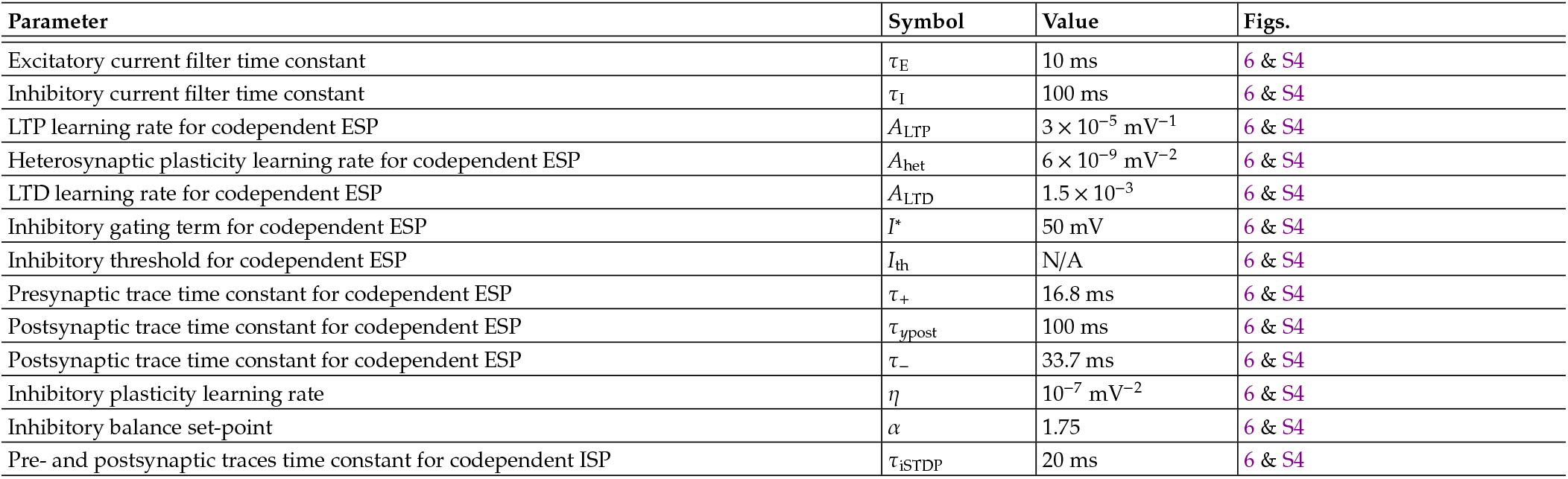
Simulation parameters for codependent synaptic plasticity model in Fig. 6 and Fig. S4.

| Parameter | Symbol | Value | Figs. |
| --- | --- | --- | --- |
| Excitatory current filter time constant | $\tau_E$ | 10 ms | 6 & S4 |
| Inhibitory current filter time constant | $\tau_I$ | 100 ms | 6 & S4 |
| LTP learning rate for codependent ESP | $A_{LTP}$ | $3 \times 10^{-5} \text{ mV}^{-1}$ | 6 & S4 |
| Heterosynaptic plasticity learning rate for codependent ESP | $A_{het}$ | $6 \times 10^{-9} \text{ mV}^{-2}$ | 6 & S4 |
| LTD learning rate for codependent ESP | $A_{LTD}$ | $1.5 \times 10^{-3}$ | 6 & S4 |
| Inhibitory gating term for codependent ESP | $I^*$ | 50 mV | 6 & S4 |
| Inhibitory threshold for codependent ESP | $I_{th}$ | N/A | 6 & S4 |
| Presynaptic trace time constant for codependent ESP | $\tau_+$ | 16.8 ms | 6 & S4 |
| Postsynaptic trace time constant for codependent ESP | $\tau_{ypost}$ | 100 ms | 6 & S4 |
| Postsynaptic trace time constant for codependent ESP | $\tau_-$ | 33.7 ms | 6 & S4 |
| Inhibitory plasticity learning rate | $\eta$ | $10^{-7} \text{ mV}^{-2}$ | 6 & S4 |
| Inhibitory balance set-point | $\alpha$ | 1.75 | 6 & S4 |
| Pre- and postsynaptic traces time constant for codependent ISP | $\tau_{iSTDP}$ | 20 ms | 6 & S4 |

**TABLE S6.** Simulation parameters for codependent synaptic plasticity model in Fig. 7, Fig. 8, and Fig. S5.

| Parameter | Symbol | Value | Figs. |
| --- | --- | --- | --- |
| Excitatory current filter time constant | $\tau_E$ | 10 ms | 7, 8, & S5 |
| Inhibitory current filter time constant | $\tau_I$ | 100 ms | 7, 8, & S5 |
| LTP learning rate for codependent ESP | $A_{LTP}$ | $3 \times 10^{-4} \text{ mV}^{-1}$ | 7, 8, & S5 |
| Heterosynaptic plasticity learning rate for codependent ESP | $A_{het}$ | $1.5 \times 10^{-8} \text{ mV}^{-2}$ | 7, 8, & S5 |
| LTD learning rate for codependent ESP | $A_{LTD}$ | $3 \times 10^{-5}$ | 7, 8, & S5 |
| Inhibitory gating term for codependent ESP | $I^*$ | 200 mV | 7, 8, & S5 |
| Presynaptic trace time constant for codependent ESP | $\tau_+$ | 16.8 ms | 7, 8, & S5 |
| Postsynaptic trace time constant for codependent ESP | $\tau_{ypost}$ | 100 ms | 7, 8, & S5 |
| Postsynaptic trace time constant for codependent ESP | $\tau_-$ | 33.7 ms | 7, 8, & S5 |
| Inhibitory plasticity learning rate | $\eta$ | $10^{-8} \text{ mV}^{-2}$ | 7, 8, & S5 |
| Inhibitory balance set-point | $\alpha$ | 1.2 | 7, 8, & S5 |
| Pre- and postsynaptic traces time constant for codependent ISP | $\tau_{iSTDP}$ | 20 ms | 7, 8, & S5 |

**TABLE S7.**
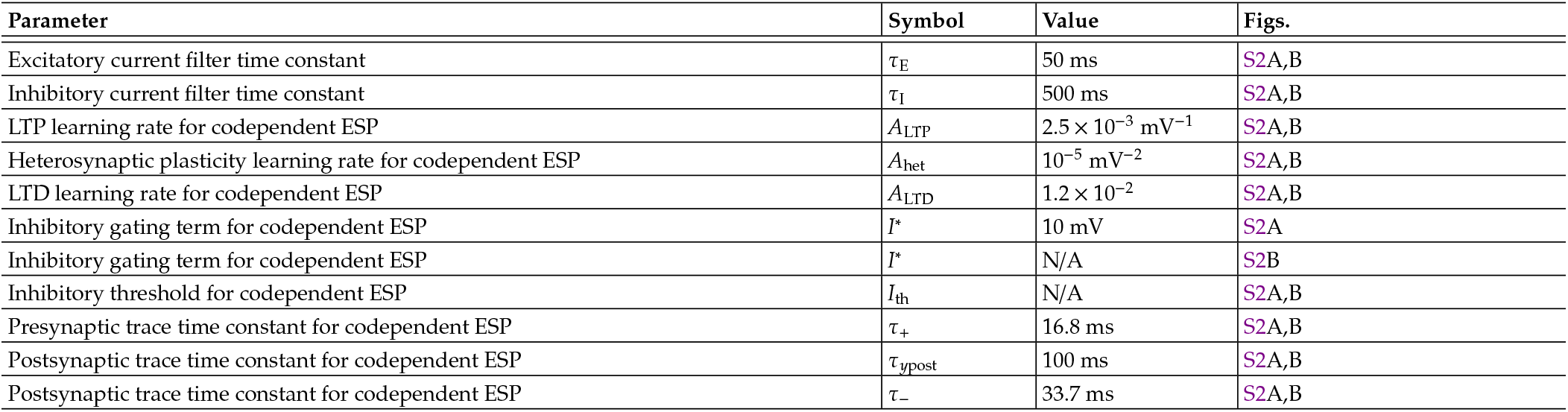
Simulation parameters for codependent synaptic plasticity model in Fig. S2A,B.

**TABLE S8.** Simulation parameters for spike- and voltage-based synaptic plasticity models in Fig. S2C,D.

| Parameter | Symbol | Value | Figs. |
| --- | --- | --- | --- |
| Neurotransmitter induced plasticity learning rate for spike-based ESP | $A_{pre}$ | 0 | S2C |
| LTP learning rate for spike-based ESP | $A_{LTP}$ | $10^{-3}$ | S2C |
| LTD learning rate for spike-based ESP | $A_{LTD}$ | $4.5 \times 10^{-2}$ | S2C |
| Presynaptic trace time constant for spike-based ESP | $\tau_+$ | 16.8 ms | S2C |
| Postsynaptic trace time constant for spike-based ESP | $\tau_y$ | 100 ms | S2C |
| Postsynaptic trace time constant for spike-based ESP | $\tau_-$ | 33.7 ms | S2C |
| LTP learning rate for voltage-based ESP | $A_{LTP}$ | $10^{-6} \text{ mV}^{-2} \text{ ms}^{-1}$ | S2D |
| LTD learning rate for voltage-based ESP | $A_{LTD}$ | $6 \times 10^{-5} \text{ mV}^{-1}$ | S2D |
| Voltage threshold for LTD | $\theta_-$ | -52 mV | S2D |
| Voltage threshold for LTP | $\theta_+$ | -55 mV | S2D |
| Presynaptic trace time constant for spike-based ESP | $\tau_x$ | 15 ms | S2D |
| Voltage trace time constant for voltage-based ESP variable $\overline{u^-}_i$ | $\tau_-$ | 10 ms | S2D |
| Voltage trace time constant for voltage-based ESP variable $\overline{u^+}_i$ | $\tau_+$ | 150 ms | S2D |

**TABLE S9.**
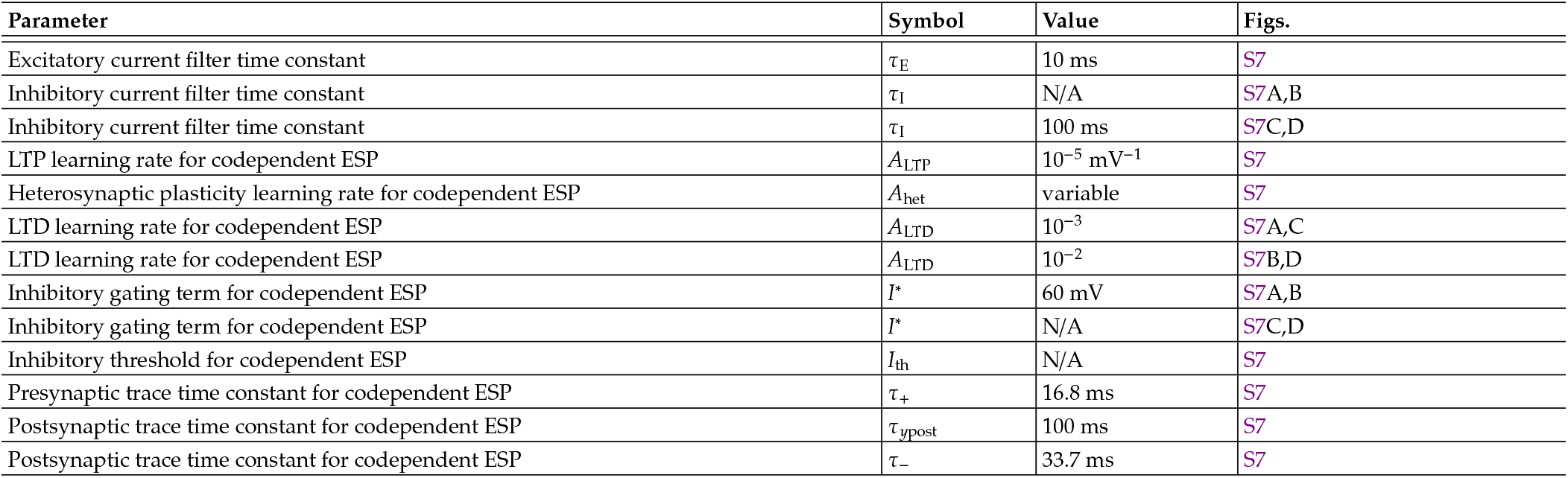
Simulation parameters for codependent synaptic plasticity model in Fig. S7.

| Parameter | Symbol | Value | Figs. |
| --- | --- | --- | --- |
| Excitatory current filter time constant | $\tau_E$ | 10 ms | S7 |
| Inhibitory current filter time constant | $\tau_I$ | N/A | S7A,B |
| Inhibitory current filter time constant | $\tau_I$ | 100 ms | S7C,D |
| LTP learning rate for codependent ESP | $A_{LTP}$ | $10^{-5} \text{ mV}^{-1}$ | S7 |
| Heterosynaptic plasticity learning rate for codependent ESP | $A_{het}$ | variable | S7 |
| LTD learning rate for codependent ESP | $A_{LTD}$ | $10^{-3}$ | S7A,C |
| LTD learning rate for codependent ESP | $A_{LTD}$ | $10^{-2}$ | S7B,D |
| Inhibitory gating term for codependent ESP | $I^*$ | 60 mV | S7A,B |
| Inhibitory gating term for codependent ESP | $I^*$ | N/A | S7C,D |
| Inhibitory threshold for codependent ESP | $I_{th}$ | N/A | S7 |
| Presynaptic trace time constant for codependent ESP | $\tau_+$ | 16.8 ms | S7 |
| Postsynaptic trace time constant for codependent ESP | $\tau_{ypost}$ | 100 ms | S7 |
| Postsynaptic trace time constant for codependent ESP | $\tau_-$ | 33.7 ms | S7 |

### Supplementary figures

**FIG. S1.**
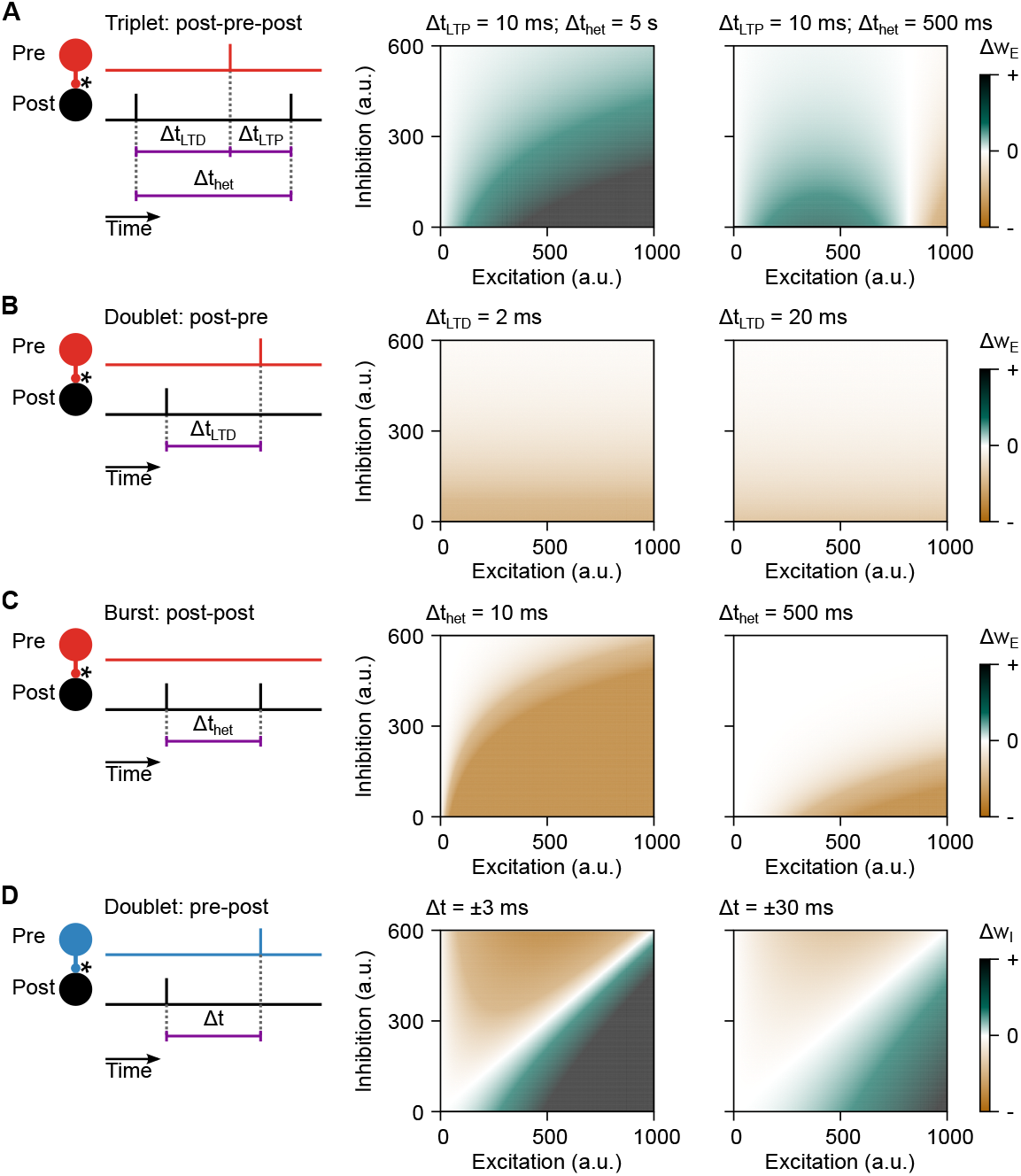
Contribution of spike times, excitation, and inhibition to weight changes for the codependent synaptic plasticity model. **A-C**, Schematics of the sequence of spikes (left), and the resulting weight change for two different spike patterns (middle and right) for codependent excitatory synaptic plasticity model as a function of the levels of excitation and inhibition during plasticity (Eq. S36). **A**, Spike triplet: post-pre-post sequence with fixed pre-before-post spike interval, Δ*t*_LTD_, and two examples for intervals between two consecutive postsynaptic spikes, Δ*t*_het_. **B**, Spike doublet: post-before-pre spike pattern with two different intervals, Δ*t*_LTD_. **C**, Postsynaptic burst with two spikes at different interspike intervals, Δ*t*_het_. **D**, Same as panel B for codependent inhibitory synaptic plasticity model (Eq. S37).

**FIG. S2.**
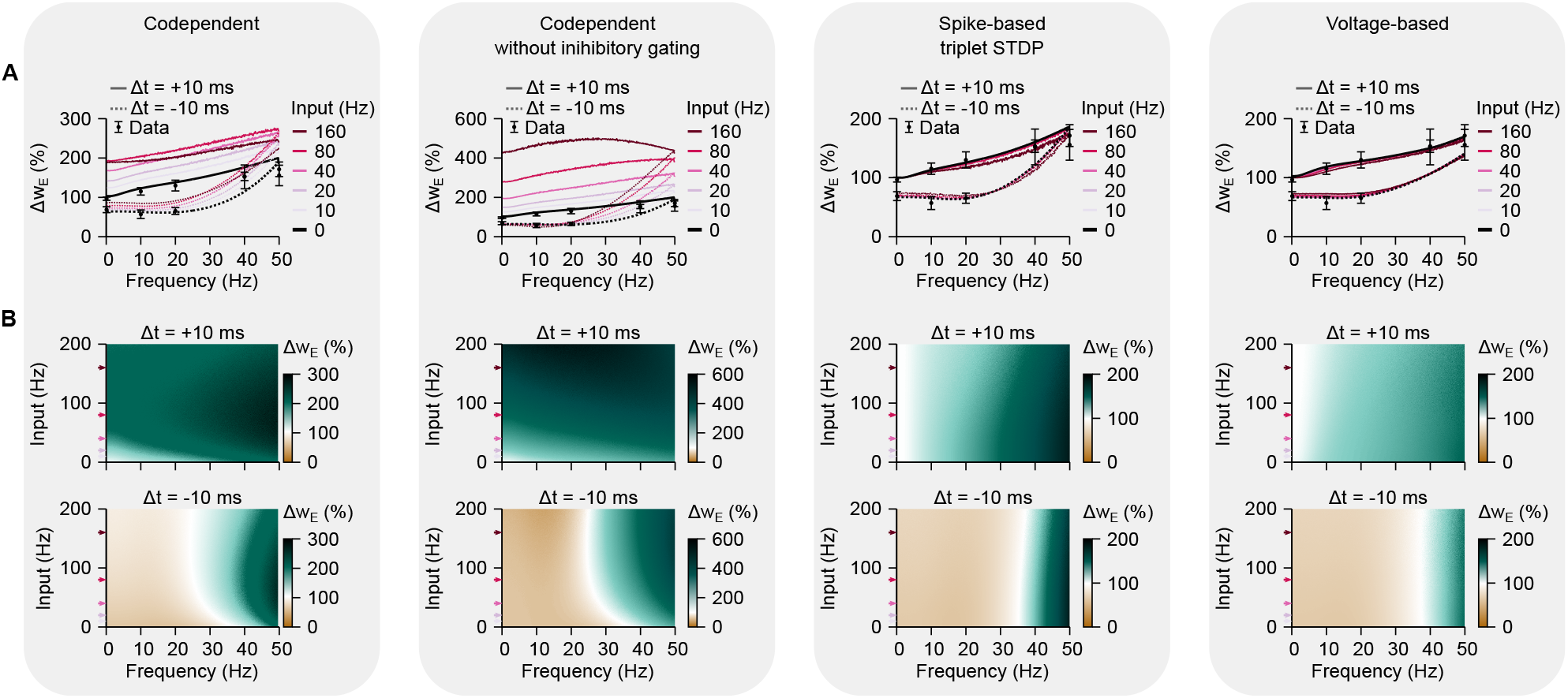
Comparison between synaptic plasticity models. **A**, Plasticity inducing protocol with pairs of pre-before-post (Δ*t* = +10 ms) and post-before-pre (Δ*t* = −10 ms) for varying spiking frequencies, and different firing-rates of neighbouring excitatory and inhibitory afferents (different colours). Plot shows changes in synaptic weight of a single connection while the other two (excitatory and inhibitory) are kept fixed. Excitatory and inhibitory weight of neighbouring synapses were chosen to keep the initial (before plasticity) excitatory and inhibitory currents balanced, and thus same average membrane potential for the same input frequency. Experimental data from Sjöström *et al*. ^17^ ; spike-based triplet spike timing-dependent-plasticity model from Pfister and Gerstner ^6^, and voltage-based plasticity model from Clopath *et al*. ^7^. Error bars indicate SEM. **B**, Weight changes as a function of input frequency (from neighbouring excitatory and inhibitory synapses; y-axis), and frequency of pairs of spikes (x-axis). Plots from the first column (‘Codependent’) are also shown in Fig. 2D,E.

**FIG. S3.**
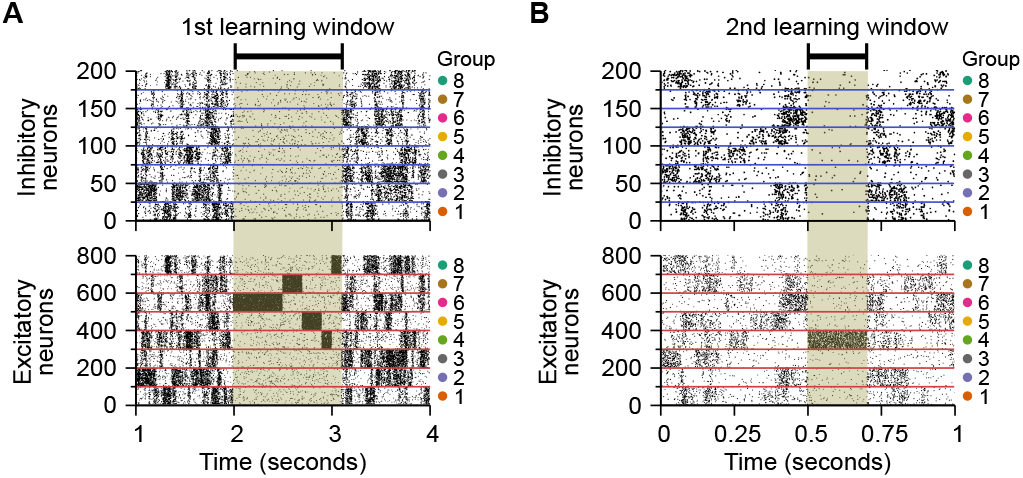
Raster plot of inhibitory (top) and excitatory (bottom) neurons used in the receptive field plasticity simulation (Fig. 4 and Fig. 5A). **A**, Input spike patterns before (*t <* 2 s), during (2 *< t <* 3.1 s), and after (*t >* 3.1 s) learning the initial receptive field profile (Fig. 4B and Fig. 5A). **B**, Sequence of input spikes for the modification of the initial receptive field profile, before (*t <* 0.5 s), during (0.5 *< t <* 0.6 s), and after (*t >* 0.6 s) the learning window (Fig. 4C).

**FIG. S4.**
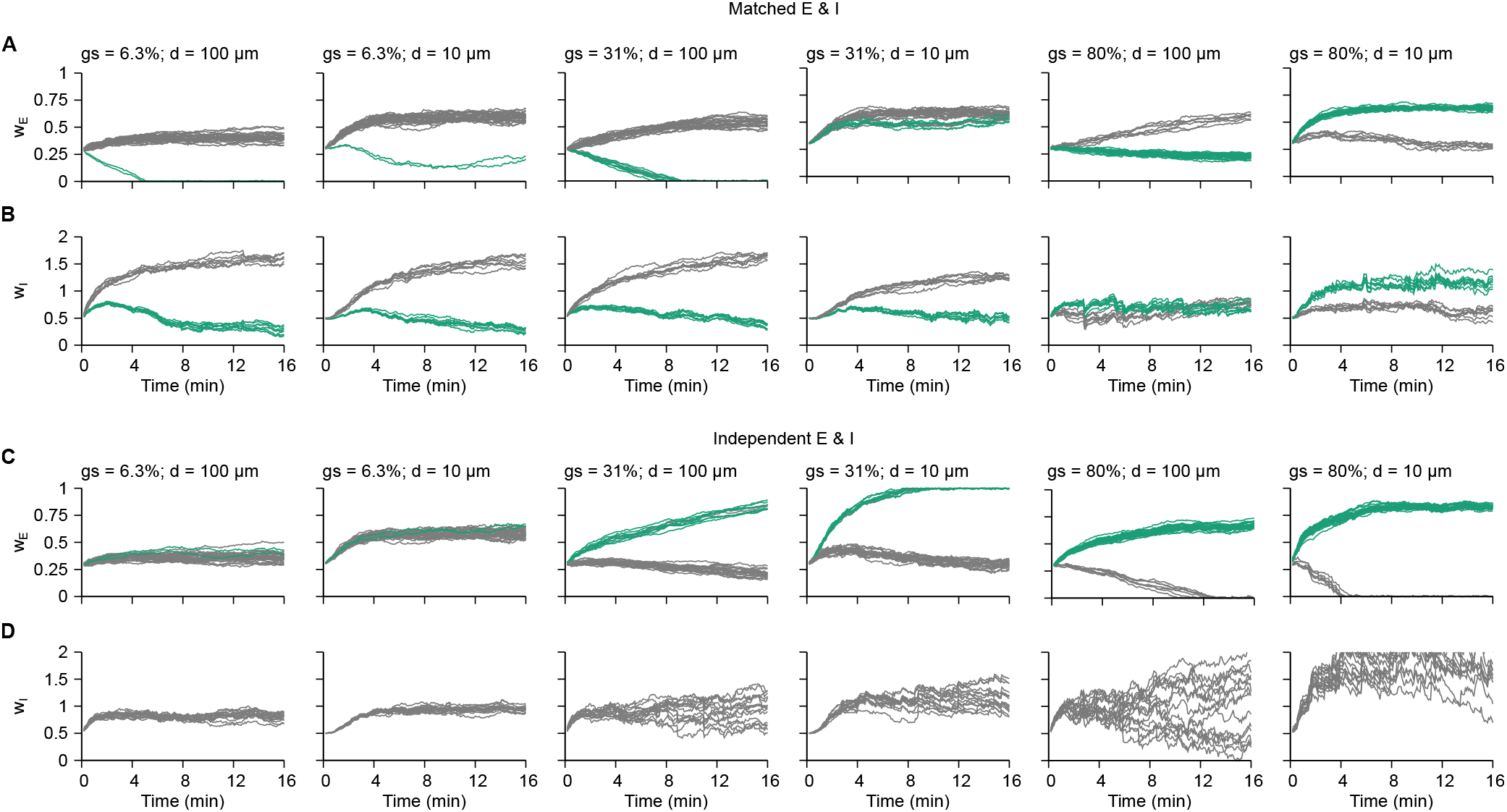
Evolution of synaptic weights connected to dendritic compartments for matched and independent E & I. **A**, Weights of co-active (green) and uncorrelated (grey) excitatory inputs with size of co-active excitatory group and distance of dendritic compartment from the soma indicated by gs and d, respectively. **B**, Weights of co-active (green; same activity pattern as co-active excitatory group) and uncorrelated (grey) inhibitory inputs. Size of co-active inhibitory group was kept fixed at half of the inhibitory population. **C**, Same as panel A, but when inhibitory inputs are independent of excitatory ones. **D**, Same as in panel B, but with no correlation between excitatory and inhibitory inputs.

**FIG. S5.**
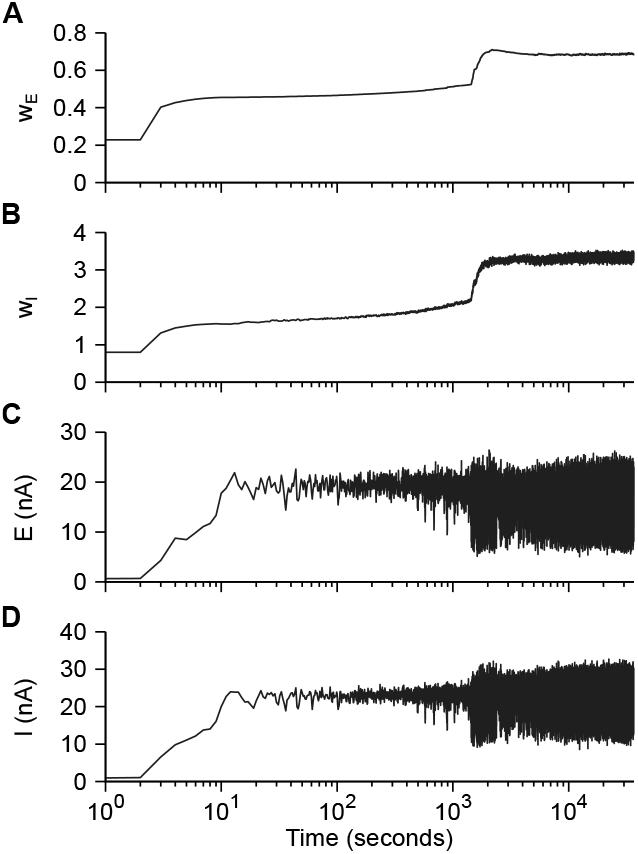
Learning period for the recurrent network (Fig. 7 and Fig. 8). **A**, Average of recurrent excitatory weights. **B**, Average of inhibitory weights onto excitatory neurons. **C**, Average of excitatory currents onto excitatory neurons. **D**, Average of inhibitory currents onto excitatory neurons.

**FIG. S6.**
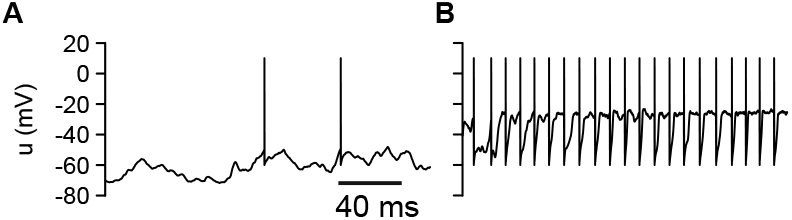
Low and high firing-rate states for neurons with adapting spiking threshold and NMDA currents. **A**, Membrane potential of a neuron in a hyperpolarised state with low firing-rate. **B**, Membrane potential of a neuron in a depolarised state with high firing-rate.

**FIG. S7.**
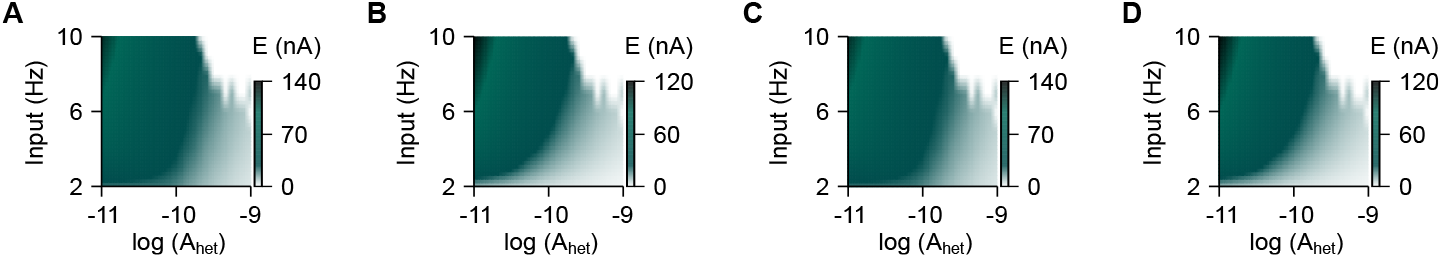
Steady state excitatory currents of a postsynaptic neuron as a function of heterosynaptic plasticity and input firing-rate. **A**, Simulation with weak LTD and weak inhibitory gating. **B**, Simulation with strong LTD and weak inhibitory gating. **C**, Simulation with weak LTD and strong inhibitory gating. **D**, Simulation with strong LTD and strong inhibitory gating. High heterosynaptic learning-rates caused vanishing of weights when inputs have higher firing-rate.

